# The miR-96 and RARγ signaling axis governs androgen signaling and prostate cancer progression

**DOI:** 10.1101/198465

**Authors:** Mark D Long, Prashant K Singh, James R Russell, Gerard Llimos, Spencer Rosario, Abbas Rizvi, Patrick R. van den Berg, Jason Kirk, Lara E Sucheston-Campbell, Dominic J Smiraglia, Moray J Campbell

## Abstract

Expression levels of retinoic acid receptor gamma (*NR1B3*/*RARG*, encodes RARγ), are commonly reduced in prostate cancer (PCa). Therefore we sought to establish the cellular and gene regulatory consequences of reduced RARγ expression, and determine RARγ regulatory mechanisms. *RARG* shRNA approaches in non-malignant (RWPE-1 and HPr1-AR) and malignant (LNCaP) prostate models revealed that reducing RARγ levels, rather than adding exogenous retinoid ligand, had the greatest impact on prostate cell viability and gene expression. ChIP-Seq defined the RARγ cistrome which was significantly enriched at active enhancers associated with AR binding sites. Reflecting a significant genomic role for RARγ to regulate androgen signaling, RARγ knockdown in HPr1-AR cells significantly regulated the magnitude of the AR transcriptome. RARγ down-regulation was explained by increased miR-96 in PCa cell and mouse models, and TCGA PCa cohorts. Biochemical approaches confirmed that miR-96 directly regulated RARγ expression and function Capture of the miR-96 targetome by biotin-miR96 identified that RARγ and a number of RARγ interacting co-factors including *TACC1* were all targeted by miR-96, and expression of these genes were prominently altered, positively and negatively, in the TCGA-PRAD cohort. Differential gene expression analyses between tumors in the TCGA-PRAD cohort with lower quartile expression levels of *RARG* and *TACC1* and upper quartile miR-96, compared to the reverse, identified a gene network including several RARγ target genes (e.g. *SOX15*) that significantly associated with worse disease free survival (hazard ratio 2.23, 95% CI 1.58 to 2.88, p=0.015). In summary, miR-96 targets a RARγ network to govern AR signaling, PCa progression and disease outcome.

**Conflict of interest:** The authors certify that they have NO affiliations with or involvement in any organization or entity with any financial interest (such as honoraria; educational grants; participation in speakers’ bureaus; membership, employment, consultancies, stock ownership, or other equity interest; and expert testimony or patent-licensing arrangements), or non-financial interest (such as personal or professional relationships, affiliations, knowledge or beliefs) in the subject matter or materials discussed in this manuscript.

**FUNDING:** *LESC* acknowledges support, in part, of Roswell Park Comprehensive Cancer Center-University of Pittsburg Cancer Institute Ovarian Cancer Specialized Program of Research Excellence National Institutes of Health [P50CA159981-01A1].

*MDL* acknowledges support of Molecular Pharmacology and Experimental Therapeutics NRSA T32 program [T32CA009072] held at Roswell Park Comprehensive Cancer Center.

*MJC* and *DJS* acknowledges support in part from the Prostate program of the Department of Defense Congressionally Directed Medical Research Programs [W81XWH-14-1-0608, W81XWH-11-2-0033] and the National Cancer Institute (NCI) grant P30CA016056 involving the use of Roswell Park Comprehensive Cancer Center’s Genomic Shared Resource.

*MJC, GL, AR, HW* and *PvdB* acknowledges support from the European Union-United States Atlantis Program [P116J090011].

*MJC* and *LESC* acknowledge support from the National Cancer Institute (NCI) grant P30CA016056 involving the use of OSUCCC The James, CCSG P30CA016058

## INTRODUCTION

Members of the nuclear hormone receptor (NR) superfamily are ubiquitously expressed across tissues and govern cell fate decisions. One NR, the androgen receptor (*NR3C4/AR*), is a key regulator of growth and differentiation in the prostate gland^37^. Genomic approaches in prostate cancer (PCa) have identified that the capacity of the AR becomes skewed with disease progression^51^. A normal component of AR signaling is to drive terminal differentiation of luminal epithelial cells and this function is disrupted in PCa^91^. The disruption, or re-wiring, of AR actions changes both the receptor sensitivity, defined as the magnitude of the transcriptional response, and capacity, defined as the selection of gene networks governed. Although the AR is a pharmacological target in later PCa stages, its disruption at early stage disease is more nuanced.

In fact, multiple NRs are expressed in the normal prostate and are disrupted in PCa. NR actions are integrated by shared genomic binding regions, shared co-factors and the co-regulation of ligand availability^61^. Non-coding RNAs including miRNA and lncRNA target NRs, their co-factors and target genes to also exert control of signaling^74, 77^. Combined, these different aspects of NR regulation ultimately modulate transcriptional sensitivity.

Previously, as a route to identify how NR networks are disrupted in cancer, we undertook an analysis of the NR superfamily across cancers in The Cancer Genome Atlas (TCGA). Our analyses revealed significantly distinct NR profiles within different tumor types^45^. Specifically, in the Taylor et al MSKCC^76^ and TCGA-PRAD PCa cohorts^13^ the retinoic acid receptor gamma (*NR1B3*/*RARG*, encodes RARγ) and glucocorticoid receptor (*NR3C1/GR*) were significantly and uniquely down-regulated. By contrast, expression of the *AR* was not significantly altered in either cohort. There was only one *RARG* mutation and relatively few CNVs detected at the *RARG* locus across these approximately 600 PCa samples.

There are three human RAR paralogs, namely RARα RARβ In PCa, RARβ appears to act as a tumor suppressor silenced by DNA methylation^10, 19^. Curiously, while there are observed roles for RARγ in prostatic development^44^, its role and regulatory functions in prostate cells and PCa remain enigmatic, as do its upstream control mechanisms. Furthermore, pharmacologic targeting of these receptors has been investigated, for example with pan and paralog specific retinoid ligands with the goal to induce differentiation^48^. However the extent to which RARs functions are directly associated with ligand activating events or indirectly through interactions with other transcription factors is similarly underexplored.

To better understand the consequences and causes of reduced RARγ expression levels in prostate cells we designed a workflow combining analyses in prostate cell lines, murine models and human tumors (Figure 1). Specifically in two non-malignant models (RWPE-1 and HPr1-AR) and in one malignant model (LNCaP) we generated two independent clones with stable RARγ knockdown. In these control and knockdown clones we examined the effects on cell viability and gene expression from either changing the baseline RARγ expression levels or adding exogenous ligand. We identified that reducing RARγ expression levels had a bigger impact on cell viability and gene expression than adding exogenous ligand. Notable in the enriched terms of the RARγ-regulated gene networks were terms related to NF-κB, hypoxia and androgen signaling. In RWPE-1 cells, we undertook RARγ ChIP-Seq to identify the RARγ cistrome. Without adding exogenous ligand RARγ significantly associated with active gene enhancers, and also significantly overlapped with the binding sites for other transcription factor functions, including AR and also the NF-κB component RELA/p65. Testing if RARγ regulated AR was undertaken by androgen-dependent transcriptomic analyses in HPr1-AR cells with stable knockdown of RARγ expression. This revealed that RARγ expression levels potently regulated the capacity and sensitivity of AR. A major regulator of RARγ expression was identified as miR-96, which is commonly elevated in PCa and associated with disease progression. MiR-96 directly bound and regulated expression of RARγand capturing the miR-96 targetome revealed that this miRNA also targeted a number of known RARγ co-factors including TACC1. Finally, tumors in the lower quartile *RARG* and *TACC1* and upper quartile miR-96 were significantly associated with aggressive PCa and disease recurrence. Together these findings suggest that RARγ expression levels potently regulate gene networks that are significantly intertwined with the regulation of AR sensitivity and capacity. Control of these actions is regulated by miR-96 and loss of this capacity predicts prostate cancer progression.

**Figure 1:**
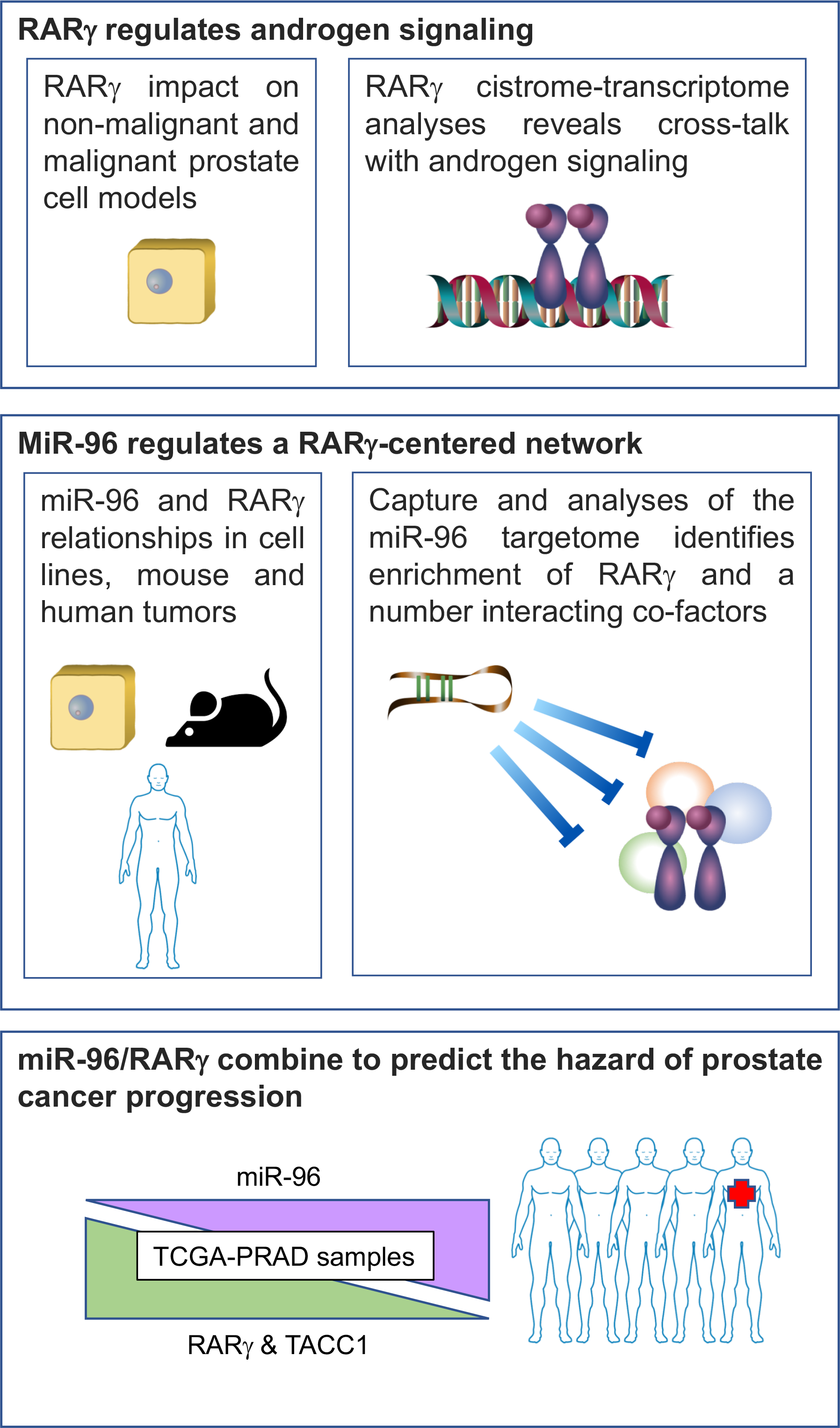
The workflow for investigating the consequences of altered RARγ expression in cell line, murine and human prostate cells, and how miR-96 regulates RARγ to drive aggressive prostate cancer.

## RESULTS

### Reduced RARγ expression in non-malignant and malignant prostate cell models increases cell viability and changes gene expression

To test the cellular impact of reduced RARγ expression levels we generated stably knocked-down RARγ levels in non-malignant prostate epithelial cells (RWPE-1) and LNCaP PCa cells using two separate RARγ targeting shRNA constructs (Figure 2A-D). The expression of the RARγ paralogs RARα and RARβ, were unaltered by the knockdown of RARγ (Figure 2A,C).

**Figure 2:**
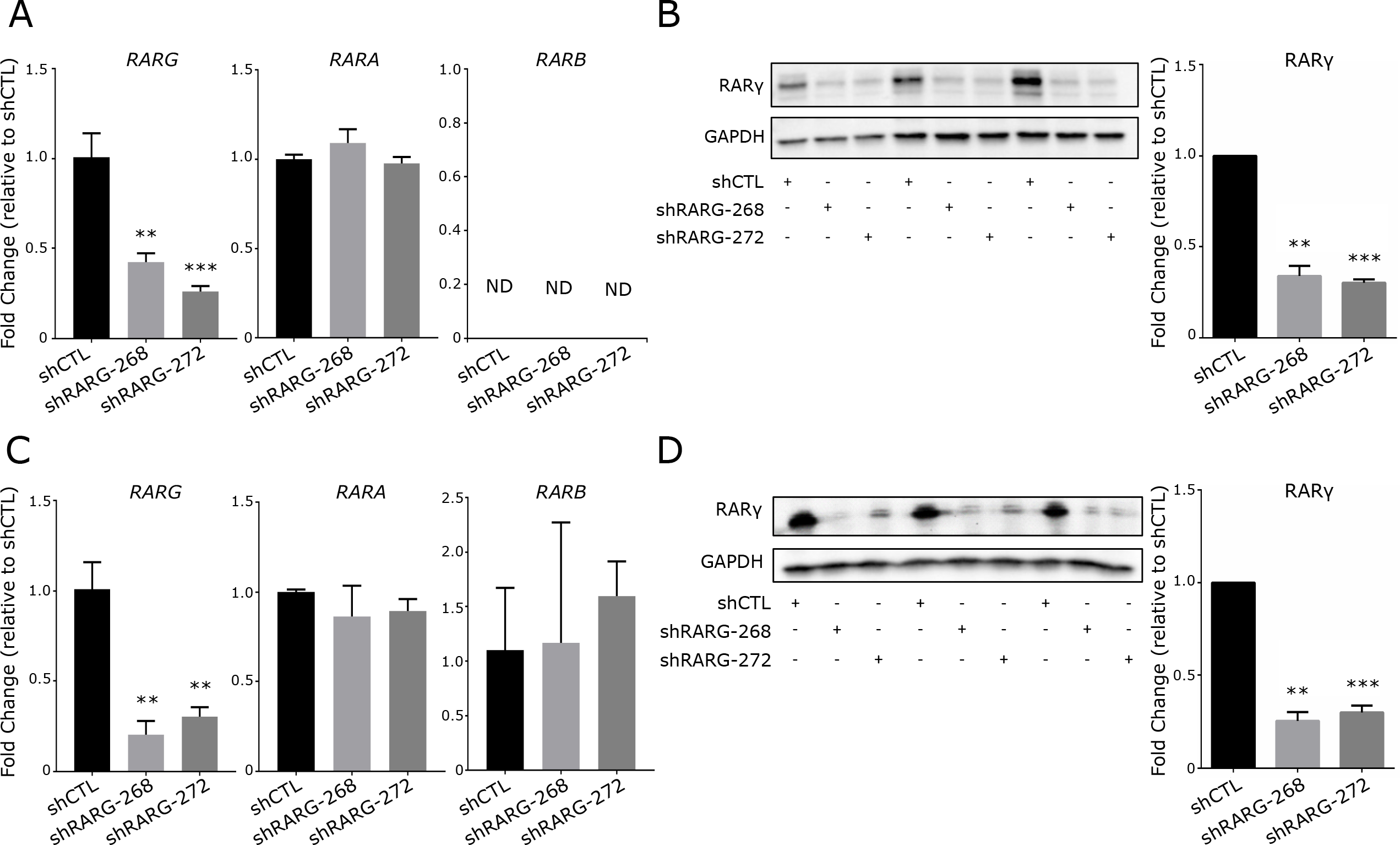
Stable knockdown of RARγ in prostate cell lines. RWPE-1 (**A,B**) and LNCaP (**C, D**) cells were each stably transfected with two shRNA constructs targeting *RARG*. (**A**) Validation of *RARG* knockdown, and unaltered expression of *RARA* and *RARB* mRNA and (**B**) RARγ protein levels in RWPE-1-shCTL and RWPE-1-shRARG cells. (**C, D**) Similar validation results are shown for LNCaP-shCTL and LNCaP-shRARG cells. Significance of difference between shRARG and shCTL cells are noted (* <0.05, ** < 0.01, *** < 0.001).

In RWPE-1 and LNCaP cells in standard cell tissue culture conditions and without exogenous retinoids, RARγ knockdown in two separate clones significantly increased viability within 96 hours (Figure 3A). By contrast, stable RARγ knockdown only slightly reduced anti-proliferative sensitivity to either all-trans retinoic acid (ATRA) or a RARγ selective ligand (CD437) in RWPE-1 cells and not at all in LNCaP cells (**Supplementary Figure 1A-D**). Also, independent of exposure to ligand, RARγ knockdown significantly reduced the G_2_/M population in both RWPE-1 and LNCaP cells, whereas reduced RARγ levels changed CD437-induced G_2_/M blockade only in RWPE-1 cells (**Supplementary Figure 1E-F**). Together these findings suggest that reducing RARγ expression has a significant impact on cell viability and cell cycle status, which is largely independent of the addition of exogenous retinoid ligand.

**Figure 3:**
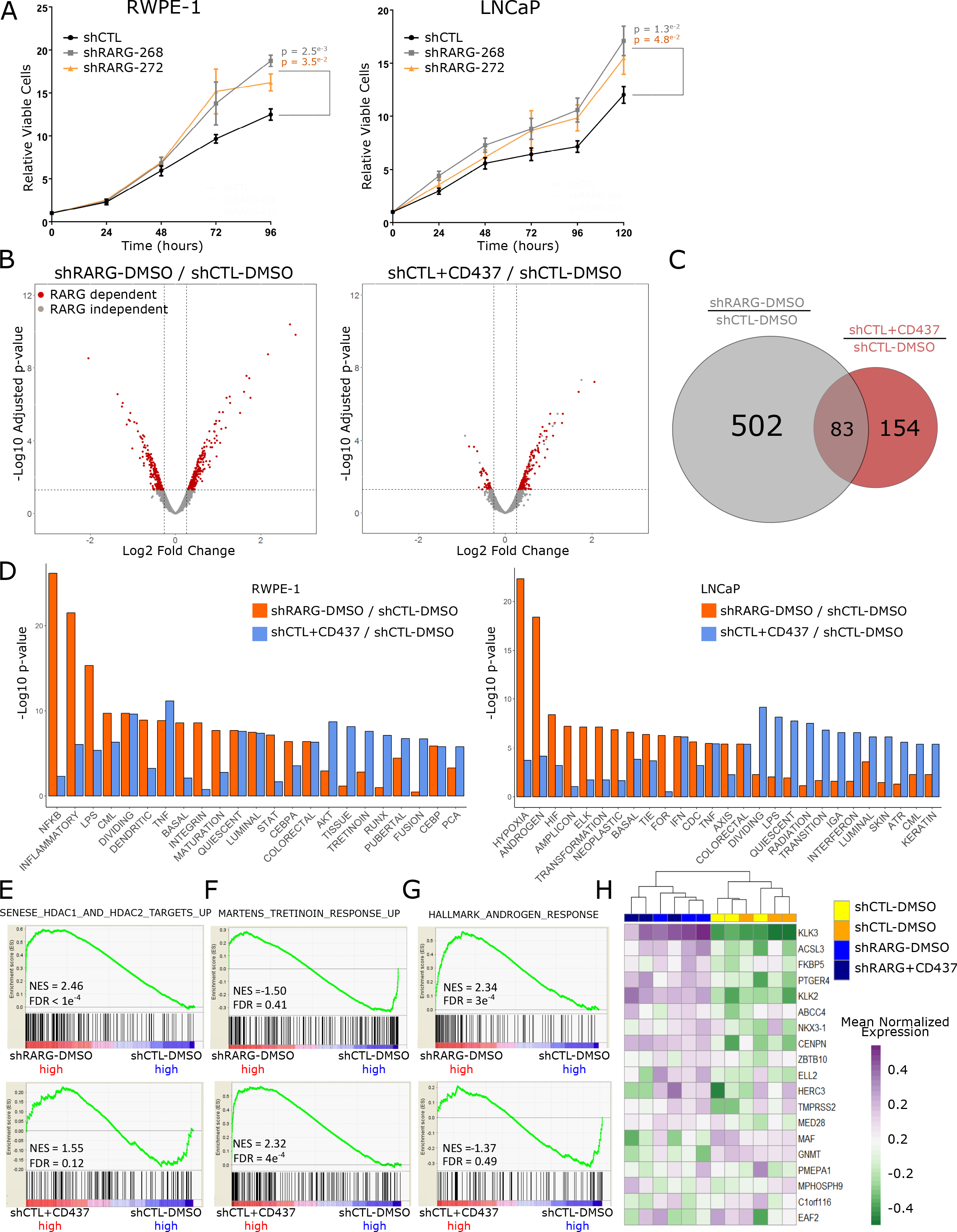
RARγ expression levels impact cell viability and gene expression in prostate cells. (**A**) Time-dependent measurements of cellular levels of ATP, as an indicator of cell viability, of each of the stable shRARG clones compared to vector controls in RWPE-1 (left) and LNCaP (right) cells. Each measurement was performed in biological triplicates, in triplicate wells. Significance of differences between viability of control and RARγ knockdown cells at the end of each study is noted. (**B**) Cells were treated in triplicate with CD437 (RWPE-1, 10 nM; LNCaP, 250 nM, 24hr) or DMSO vehicle control and gene expression was profiled using Illumina microarray (Illumina HT12v4) using DESeq2. Volcano plots depicting expression changes upon RARγ knockdown or in response to exogenous RARγ specific ligand (CD437) in RWPE-1 cells. Dotted lines indicate DEG thresholds (p.adj < 0.05, fold change of +/− 1.2), and red dots represent RARγ dependent DEGs. Genes regulated by exogenous ligand were calculated as those that were differentially expressed in shCTL cells treated with ligand (shCTL+CD437/shCTL-DMSO) and not in shRARG cells treated with CD437. (**C**) Venn diagram depicting number of determined DEGs associated with reducing RARγ expression levels, and those from adding exogenous ligand. (**D**) Summary of significantly enriched pathways from gene set enrichment analyses (GSEA) (NES > 1.8, FDR q.val <0.05) associated with reducing RARγ expression levels, and those from adding exogenous ligand in RWPE-1 (left) and LNCaP (right) cells. The top enriched meta-groups amongst significant GSEA sets are indicated. (**E,F**) Examples of top significant GSEA pathway enrichments observed in RWPE-1 cells and (**G**) in LNCaP cells. (**H**) Heatmap depicting the relative expression of a panel of androgen response genes defined previously by the TCGA consortium to reflect AR signaling^14^. Unsupervised hierarchical clustering of the gene expression patterns of these AR-regulated genes separated shRARG from shCTL cells.

Stably reducing RARγ expression levels also exerted a significant impact on gene expression, compared to non-targeting vector controls. Meanwhile the RARγ dependent CD437-induced response, defined as genes differentially regulated by CD437 in shCTL cells but not (or to a significantly lesser extent) in shRARG cells, was more modest (RWPE-1) or negligible (LNCaP) (Figure 3B-C, **Supplementary Figure 2A-D**). For instance in RWPE-1 cells, 605 differentially expressed genes (DEGs) (1.2 fold change, Benjamini-Hochberg adjusted p-value < 0.05) were associated with the knockdown of RARγ regulation, whereas only 237 genes were responsive to CD437 (10 nM, 24 hr) in a RARγ dependent manner.

To infer gene functionality resulting from changes in RARγ expression, Gene Set Enrichment Analysis (GSEA) was applied to the RARγ dependent transcriptomes. In RWPE-1 cells, 435 pathways were enriched (normalized enrichment score (NES) > 1.8, FDR q-val < 0.05) in at least one comparison (**Supplementary Figure 2E**). Amongst these, 196 were associated uniquely with stable knockdown of RARγ and 201 with the addition of CD437, while 38 were common between categories.

Frequency mining of common GSEA pathway terms combined with hypergeometric testing established largely distinct enrichment of meta-groups (e.g. terms associated with NF-κB) between stable RARγ knockdown or CD437 treated conditions (Figure 3D). In RWPE-1 cells, the RARγ stable knockdown transcriptome was significantly enriched for terms associated with NF-κB and histone deacetylase (HDAC) function (Figure 3E) whereas the CD437 treated transcriptome was enriched for terms related to Tretinoin (the commercial name for ATRA) (Figure 3F), as well as for ERα function. In LNCaP cells the stable knockdown of RARγ gene regulatory effects were most pronounced (**Supplementary Figure 2C-D**) and highly enriched for hypoxia and AR responses (Figure 3G, **Supplementary Figure 2F**).

We further explored the enrichment of AR terms in the transcriptome of stable knockdown of RARγ in LNCaP. Specifically, we used a previously compiled AR target gene panel ^13^ to demonstrate that expression of these genes alone significantly distinguished LNCaP-shRARG cells from controls, in a CD437-independent manner (Figure 3H). That is applying unsupervised hierarchical clustering to expression patterns of these genes distinguished the LNCaP-shRARG from CD437 treated clones. Furthermore, the expression of *KLK3* and *TMPRSS2* were altered upon knockdown of RARγ in LNCaP cells and its loss altered DHT regulation of these genes (**Supplementary Figure 2G**).

In total, stable knockdown of RARγ revealed functions of this receptor that are substantial and intertwined with other transcription factors, notably the AR and NF-κB, and these actions appear to be largely independent of the impact of treating cells with retinoid ligand.

### The RAR_γ_ cistrome significantly overlaps with active enhancers, AR binding and associates with aggressive PCa

Having established that changing RARγ expression levels can have a significant impact on gene expression, even in the absence of exogenous ligand, we next wished to measure the genomic distribution of RARγ in the presence and absence of exogenous retinoid ligand. To avoid the Type I error rate due to RARγ paralog noise we used a GFP tagged RARγ contained in a BAC clone (kind gift of Dr. Kevin White (University of Chicago)). Indeed these investigators demonstrated that commercially available RARγ antibodies were not able to yield high quality chromatin for ChIP studies^30, 39^. Although GFP-RARγ was modestly elevated compared to endogenous RARγ this does not appear to be supra-physiological, like that which may occur with CMV and other strong regulatory promoters. Therefore RARγ-EGFP was stably transfected in RWPE-1 cells^30^; optimization studies revealed significant enrichment at previously established RARγ dependent genes including SMAD3^17, 67^ and FOXA1^18^ (**Supplementary Figure 4A-C**).

**Figure 4:**
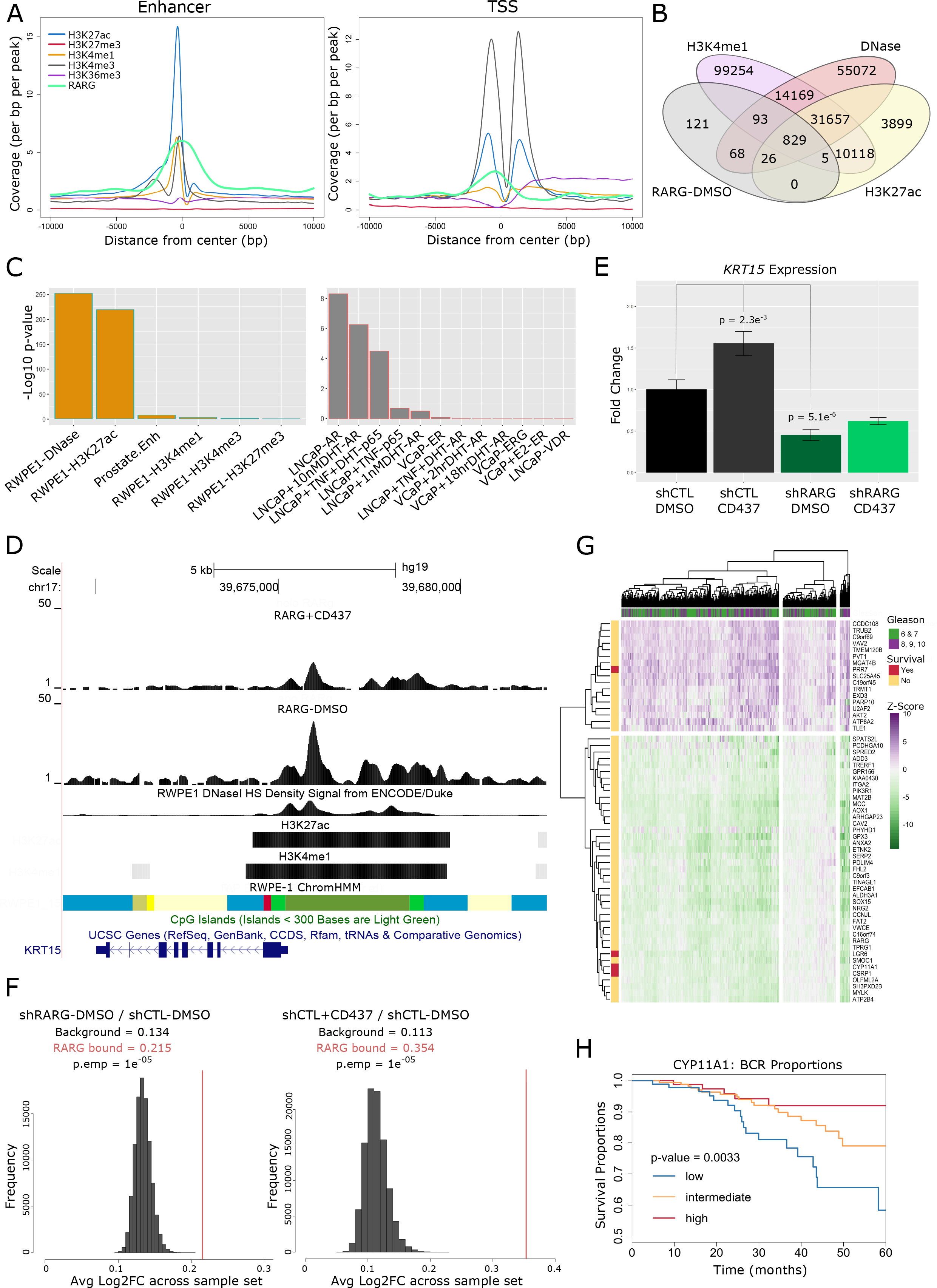
The RARγ cistrome is enriched at active enhancers of genes that are also enriched for androgen receptor binding and associate with aggressive prostate cancer. Stable transfection of a bacterial artificial chromosome containing a fusion *RARG-EGFP* gene transcribed from the endogenous *RARG* promoter was undertaken to generate RWPE-1-RARG-EGFP clones. Stably expressing RARγ-EGFP RWPE-1 cells were treated in triplicate with either CD437 (10 nM, 2 hours) or vehicle (DMSO), and subjected to ChIP-Seq undertaken with an EGFP antibody. (**A**) Significant RARγChIP-seq peaks (p.adj <0.1) were determined with Rsubread and csaw. Cross profiling of significant RARγChIP-seq peaks and the indicated histone modifications also from RWPE-1 (GSE63094) at genomic loci performed using annotatePeaks available from the HOMER (Hypergeometric Optimization of Motif EnRichment) suite. Genomic profiles are centered at enhancer regions (Overlapping of H3K27ac and H3K4me1) (left), and TSS loci of all expressed genes in RWPE-1 (right). (**B**) Overlap of unstimulated RARγ binding peaks with select publicly available epigenetic datasets from RWPE-1 cells was determined with ChIPpeakAnno using a maximum gap of 500 bp; H3K4me1 and H3K27ac (GSE63094) and DNase sensitivity (GSM1541008). (**C**) The negative log10(pValues) of the overlaps between data sets determined with ChIPpeakAnno were visualized for RARγ binding peaks and a wider panel of publicly available data; prostate enhancers (FANTOM), LNCaP AR (GSE48308), VCaP AR (GSE84432), ERα (GSE43985) and LNCaP NF-κB (GSE83860) cistromes. (**D**) Representative RARγbinding site upstream of the *KRT15* TSS showing coincident binding with H3K4me1 and H3K27ac in RWPE-1 cells. The ChromHMM (grass green = low transcription; bright green = Active enhancer; red = Transcriptional enhancer; blue = Active TSS; pale yellow = Poised transcription; yellow = Poised enhancer), CpG island and DNase sensitivity tracks are also shown. (**E**) The impact of RARγ knockdown on *KRT15* expression from expression profiling in RWPE-1 cells by microarray (Figure 1). (**F**) Bootstrapping approach (using boot) to test the statistical strength of relationships between genes annotated to be bound by RARγ (+/− 7.5 kb from the TSS) and those modulated by RARγ knockdown (left) or in the presence of CD437 ligand (right) in RWPE-1 cells. The red line is the mean observed fold change for the indicated gene set derived from the microarray data (Figure 1) and the distribution is simulated data from the same data-set. The significance between the observed and simulated expression of the same number of genes is indicated (**G**) Expression heatmap of annotated RARγ cistrome genes in the TCGA-PRAD cohort (pheatmap). Genes were filtered to identify those that were altered by more than +/−2 Z-scores in 35% of tumors relative to normal samples. Unsupervised hierarchical clustering grouped tumors that significantly distinguished higher Gleason Grade (Gleason > 7), after adjusting for age (Pearson’s Chi-squared test X-squared = 51.187, df = 35). For each gene, Kaplan-Meier curves (survival) were generated as time to tumor recurrence, indicated by biochemical progression, and those genes which identify significantly (p.adj <0.1) shorter disease free survival are indicated. (**H**). Illustrative Kaplan-Meier curve for *CYP11A1* expression depicting the significantly reduced time to 5 year biochemical recurrence post radical prostatectomy.

In the absence of exogenous retinoid, reflecting the transcriptomic data, the RARγ cistrome was considerably larger (1256 peaks (p.adj < 0.1)) than the CD437-dependent cistrome of 360 peaks, of which 316 were shared and 44 were unique. Motif analyses of RARγ cistrome in the absence of exogenous retinoid revealed significant enrichment of transcription factor (TF) motifs for members of the AP1 family (FOSB/JUN; E-value 1.2e^−80^), the homeobox family (OTX2, PITX1; E-value 9.2e^−12^) and NRs (RXRs, RARs; E-value 2.4e^−9^). By contrast, after the addition of CD437 (10 nM, 2 hour) the RARγ cistrome was enriched uniquely for other homeobox family members (ONEC2, HXC10; E-value 1.5e^−14^), and PAX5 (E-value 6.1e^−7^).

We characterized the RARγ cistrome further by examining its overlaps with publicly available ChIP-Seq data sets. We focused on two sources; firstly ChIP-Seq and other genomic data sets derived in normal prostate cells and tissue, and secondly those from across PCa models for the AR and other nuclear receptors, as well as NF-κB and other PCa relevant factors including ERG. Therefore, we examined the RARγ cistrome genomic distribution and overlap, within 100bp, with ChIP-Seq data available in RWPE-1 cells, namely DNAse sensitivity, histone modifications (H3K27ac, H3K27me3, H3K4me1, H3K4me3) (GSE63094, GSM1541008), and the chromatin states track derived with ChromHMM^22^ (personal communication, Dr. Mathieu Lupien (University of Toronto)). Alongside these, we examined the overlap of the RARγ cistrome with normal prostate epithelial cell enhancers derived from the FANTOM consortium^4^. Secondly, the overlap with other nuclear receptors (AR, VDR, ER, VDR), NF-κB^51^ in LNCaP cells was examined as well as AR and ERα in VCaP (Figure 4A-C).

In the first instance, we used read density mapping and hypergeometric testing to examine the extent and significance of the overlaps between RARγ and other cistromes. In the absence of exogenous retinoid ligand, the RARγ cistrome significantly overlapped with regions in RWPE-1 cells that were occupied with H3K27ac, H3K4me1 marks associated with enhancer status (Figure 4A, **left**). Additionally, substantial enrichment of RARγ binding was also observed at regions upstream yet proximal of expressed TSS loci associated with high levels of H3K4me3 (Figure 4A, **right**). Specifically, 829/1256 RARγ binding sites overlapped with the combined H3K27ac, H3K4me1 and DNase cistromes, also from RWPE-1, suggesting that in the absence of exogenous retinoid ligand RARγ is commonly found in open chromatin active enhancer regions; the significance of overlaps is shown (Figure 4B,C). Reflecting the RARγ transcriptomic data, the RARγ cistrome significantly overlapped with LNCaP AR data sets ^85^, and DHT-dependent and TNF-stimulated p65 cistrome ^51^ (Figure 4C). By contrast, there was no significant overlap between the RARγ cistrome and either AR, VDR or ERG binding in the TMPRSS2-ERG fusion positive VCaP cell line.

RARγ binding at active enhancer regions was also supported by ChromHMM states derived in RWPE-1 cells showing binding of RARγ at sites flanking transcription sites. An example of this is shown for the *KRT15* gene where the location of the binding site is unambiguously related to the gene (Figure 4D).

Next, we sought to dissect transcriptome-cistrome relationships. In the first instance candidate level-relationships were tested. For example, in the absence of exogenous retinoid ligand expression of *KRT15* was both significantly reduced by the knockdown of RARγ (p = 5.1e^−6^) and induced by CD437 (p = 2.3e^−3^) (Figure 4E). To test the genome-wide level of significance transcriptome-cistrome relationships, both in the absence and presence of exogenous retinoid ligand, RARγ cistromes were annotated to within 7.5 kb of known gene TSS locations. The absolute magnitude of differential expression upon stable RARγ knockdown of these cistrome proximal genes was compared with that of all expressed genes in RWPE-1 cells (Figure 4F, **left**). To test how the associations between RARγ binding and RARγ dependent change in gene expression we used a random sampling with replacement, or bootstrapping approach, as developed previously^46^ (**Supplementary Table 1**). This approach revealed that RARγ cistrome associated genes were significantly more differentially regulated upon knockdown of RARγ than would be predicted by chance (p.emp = 1e^−5^) supporting a functional relationship between RARγ binding and expression control. A similar observation was made regarding CD437 regulated expression and CD437 dependent RARγ cistrome associated genes (p.emp = 1e^−5^) (Figure 4F, **right**).

Next, we tested the expression relationships of RARγ and the RARγ-annotated genes in the TCGA-PRAD and MSKCC cohorts. We used a Kolmogorov–Smirnov test (K–S test) to examine whether the distribution of correlations between expression of *RARG* with experimentally derived RARγ-annotated genes deviated significantly from that of its correlation with all genes. In the TCGA-PRAD and MSKCC cohorts the correlation between RARγ and RARγ-annotated genes was significantly more positive than predicted by chance **(Supplementary Figure 4A**), again suggesting functional relationships between RARγ and its target genes.

Next, we filtered RARγ-annotated genes (altered by greater than +/−2 Z-scores in more 35% tumors relative to normal tissue) and identified 58 genes. Unsupervised hierarchical clustering of the gene expression patterns identified distinct clusters of patients (Figure 4G) and distinguished a cluster of high Gleason grade tumors (after adjusting for age; p-value = 0.038). We also reasoned that higher Gleason grade tumors were associated with worse outcome and therefore we tested relationships between the expression of individual genes and disease free survival. After FDR correction, four of these genes were individually significantly associated with disease free survival including; the steroidogenic enzyme *CYP11A*^9^ (Figure 4H); *LGR6*, a WNT regulator^41^ and implicated in breast cancer; *PRR7*, a central regulator of CLOCK.

### Reduced RARγexpression alters AR signaling capacity and sensitivity

Combined, the RARγ knockdown phenotype, transcriptome and cistrome data support the concept that in the absence of exogenous retinoid ligand RARγ functions are substantially intertwined with other TFs including the AR. Therefore, we sought to test the genome-wide impact of RARγ expression on AR function directly in non-malignant prostate epithelial HPr1-AR cells.

Again, generating stable RARγ knockdown reduced RARγ levels 60-80% while RARα and RARβ remained unaffected, which was not altered by DHT treatment (**Supplementary Figure 5A-E**). HPr1-AR cells constitutively express AR, and undergo androgen-induced differentiation from a basal-like towards a more luminal-like phenotype concomitant with inhibition of proliferation^16^. Stable RARγ knockdown changed the anti-proliferative response induced by DHT, which was significantly dampened in RARγ knockdown HPr1-AR cells (Figure 5A).

**Figure 5:**
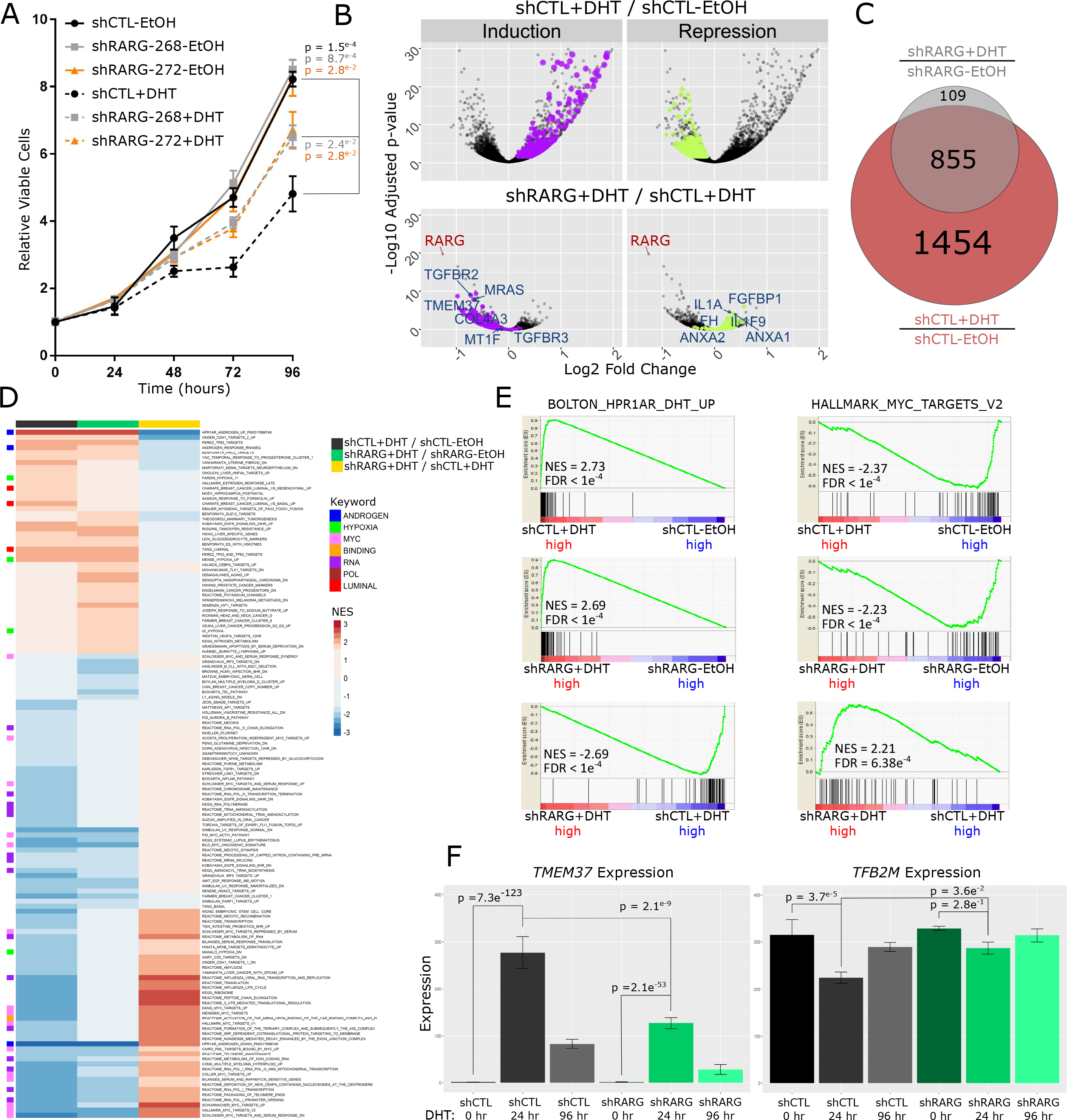
RARγ governs androgen induced responses in non-malignant HPr1-AR prostate epithelial cells. HPr1-AR cells were stable transfected with shRNA to RARG and the response to DHT measured. (**A**) Cell viability of HPr1-AR-shCTL and HPr1-AR-shRARG cells in the absence or presence of 10 nM DHT was measured for up to 96 hours. Significance of differences between triplicate experiments between viability of indicated experimental groups at the end of the study is noted. (**B**) HPr1-AR-shCTL and HPr1-AR-shRARG cells were treated in triplicate with DHT or vehicle control for 24 hours and gene expression profiled via RNA-seq and analyzed with Rsubread/DESeq2. Volcano plot depicting DHT-induced gene expression changes in HPr1-AR-shCTL cells (top). Highlighted genes represent those that display induction (purple) or repression (green) in response to DHT in HPr1-AR-shCTL expression, and which was significantly dampened in HPr1-AR-shRARG cells (bottom); 6 genes are labelled that had the greatest magnitude of dampened response, including a loss in induction of *TMEM37* and a loss of repression of *FGFBP1* (**C**) Venn diagram representing the number of DEGs determined after DHT treatment in HPr1-AR-shCTL and HPr1-AR-shRARG cells. Specifically, 1454 of the 2309 (63.0%, hypergeometric p-value < 2e^−16^) DHT-regulated DEGs were either significantly less induced, or had significantly reduced repression in shRARG cells. (**D**) Heatmap depicting normalized enrichment scores (NES) of all enriched pathways related to DHT treatment in different comparisons (NES > 1.8, FDR q-value <0.05). Select meta-groups from keyword enrichment analysis are depicted. (**E**) Example of top significantly upregulated (left) and downregulated (right) GSEA pathways upon DHT treatment. Enrichments are shown for each comparison, as well as (**F**) an expression profile of a representative gene from each gene set (left, *TMEM37*, right *TFB2M*).

We therefore investigated whether, in the absence of exogenous retinoid ligand, RARγ functions govern AR signaling globally through RNA-Seq profiling of RARγ knockdown HPr1-AR cells treated with DHT. Analyses confirmed strong reduction in *RARG* levels, being the most significantly repressed gene when comparing RARγ knockdown HPr1-AR to control cells (**Supplementary Figure 5E, Figure 5B). Indeed at the transcriptome level, RARγ knockdown substantially reduced the sensitivity of the DHT-dependent transcriptome at both 24 and 96 hours of exposure (Figure 5B-C, Supplementary Figure 6A-C**). For instance tested the mean of the log fold change between the two states and revealed that at 24 hours, 1454 of the 2309 (63.0%) DHT-regulated DEGs were either significantly less induced (Figure 5B, purple) or significantly less repressed (Figure 5B, green, p.adj<2e-16). Amongst those genes with the greatest shift in sensitivity included several established AR targets and AR pathway interacting genes including *TMEM37*, *MRAS*, *IL1A* and *ANXA1*. This shift in sensitivity was almost uniformly directional, as in almost all cases there was a dampening effect for both induced and repressed genes.

**Figure 6:**
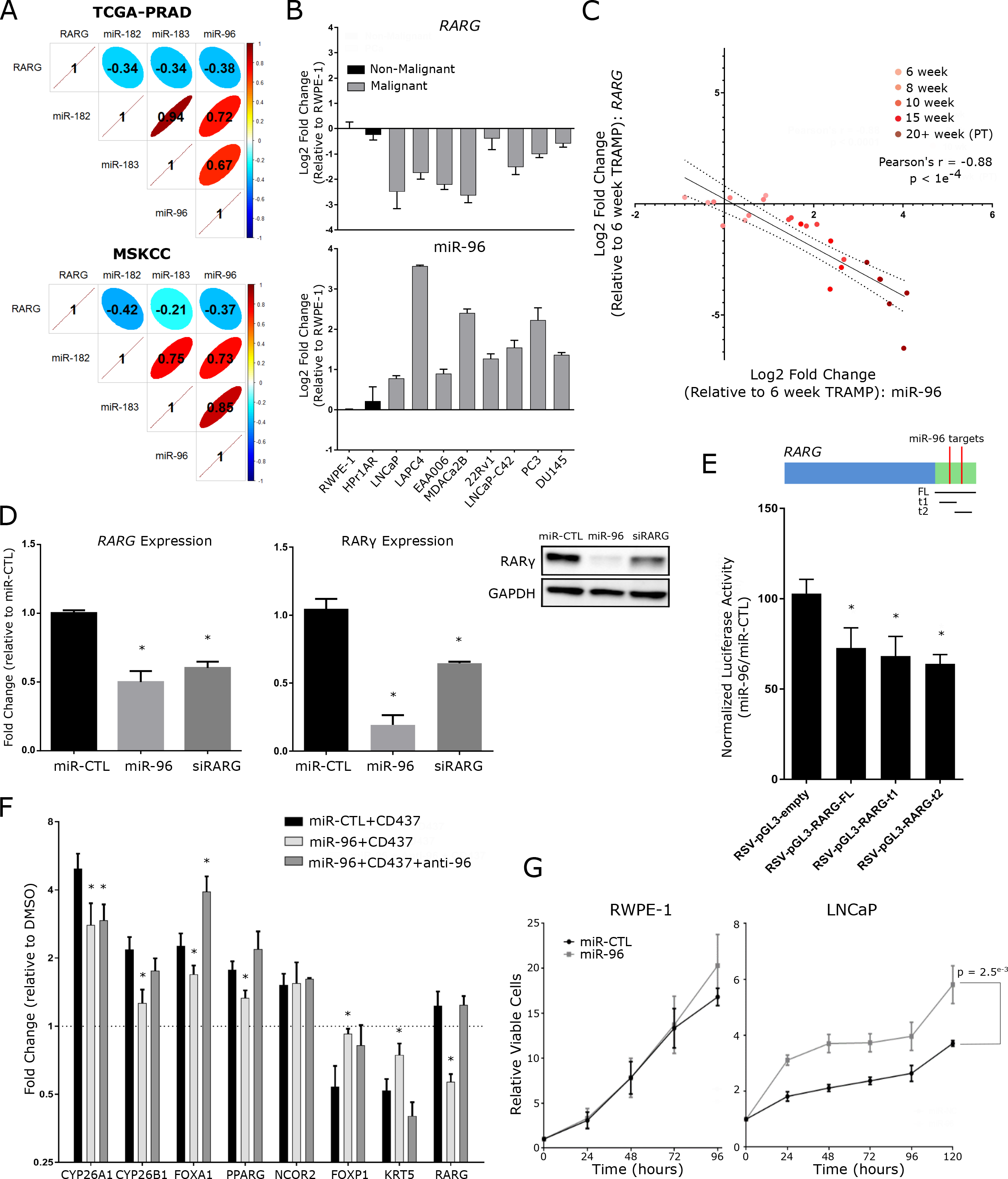
MicroRNA-96 directly targets and regulates *RARG* in prostate cells. (**A**) Cross-correlation matrices depicting the relationships between *RARG* and miR-96 cluster member expression in PCa samples from MSKCC and TCGA-PRAD cohorts (corrplot). (**B**) Relative expression of miR-96 (bottom) and *RARG* (top) across 10 prostate cell lines representing different stages of PCa progression. The cell models examined comprised immortalized RWPE-1 and HPr1AR non-malignant prostate epithelial cells, LNCaP, LAPC4 and EAA006 androgen sensitive PCa cells, MDAPCa2b, 22Rv1 and LNCaP-C42 CRPC cells, as well as PC3 and DU145 cells derived from distant metastases. (**C**) Correlation analyses of *RARG* with miR-96 expression over the course of palpable tumor (PT) development in TRAMP. The strength and significance of the correlation is indicated. Tumors from 5 mice were examined at each time point and compared to the mean of 10 6-week old wild type mice; 2 6-week tumors were dropped due to technical failure to generate high quality RNA (**D**) *RARG* mRNA (left) and RARγ protein expression (right) in RWPE-1 cells after 48 hour transfections with miR-96 mimics or siRNA targeting *RARG*. Significance of difference between targeting and control cells are noted (* <0.05). (**E**) Luciferase assay assessing direct targeting of miR-96 to the full length (FL) RARG 3’UTR or individual predicted target sites (t1, t2) within the RARG 3’UTR. Either miR-96 or miR-CTL mimics (30 nM) were transfected into RWPE-1 cells for 48 hours along with indicated RSV-pGL3 constructs and pRL-CMV Renilla luciferase expressing vectors, and luciferase activity measured by Dual-Glo Luciferase Assay System in triplicate. Significance of difference between luciferase construct with and without *RARG* 3’UTR sequences cells are noted (* < 0.05). (**F**) RWPE-1 cells were pretreated with miR-CTL, miR-96 mimic (30 nM), or combination of miR-96 mimic and antagomiR-96 for 48 hours prior to CD437 exposure (10 nM) for 24 hours, and candidate transcripts measured by RT-qPCR in triplicate. Induction relative to untreated control for each condition is shown, and significance of CD437 induction/repression between miRCTL and miR-96 groups are indicated (* < 0.05). (**G**) Cell viability of RWPE-1 (left) and LNCaP (right) cells for up to 120 hours post-transfection with either miR-96 or nontargeting control (NC) mimics was measured in triplicate. Significance of differences between viability of indicated experimental groups at the end of the study is noted.

For example, *TMEM37* is strongly induced by DHT, and RARγ loss significantly dampens this induction (Figure 5F), whereas *ANXA1* was repressed by DHT, but to a lesser extent upon RARγ loss.

To this point, the switch in the sensitivity response to RARγ loss was generally a reduction of DHT-regulation. However, the capacity of DHT regulation was also shifted, albeit to a lesser extent, as a small number of genes (109) gained unique DHT-regulation following RARγ knockdown in HPr1AR cells at 24 hours (Figure 5C). Thus, RARγ knockdown substantially impacts the sensitivity of the AR-regulated transcriptome and appears to have a more modest impact on the capacity of the AR-regulated transcriptome.

GSEA on the AR-regulated transcriptome in control and RARγ knockdown cells revealed pathways that were selectively disrupted by RARγ knockdown (Figure 5D,E). High enrichment for androgen response, MYC and hypoxia associated pathways^16^ were clear upon DHT exposure in control cells and are consistent with a shift towards luminal differentiation and an anti-proliferative response to DHT. In RARγ knockdown cells the extent of AR pathway regulation, both induction and repression events, by DHT was significantly reduced (Figure 5D). For example, an established HPr1-AR androgen response pathway (Figure 5E, **left**) was highly elevated in DHT treated relative to untreated control cells (NES = 2.73, upper panel), and while still elevated in treated RARγ knockdown cells (NES = 2.69, middle panel), the extent of this induction was substantially dampened (NES = −2.69, lower panel). Conversely, repression of MYC regulated pathways (Figure 5E, **right**) was apparent in treated control cells (NES = −2.37, upper panel) and while still apparent in RARγ knockdown cells (NES = −2.23, middle panel), the extent of repression was also reduced (NES = 2.21, lower panel). Interestingly, a prominent RARγ binding peak was identified in an enhancer region upstream of MYC (**Supplementary Figure 6D**). Candidate examples from androgen response (*TMEM37*) and MYC associated (*TFB2M*) pathways are shown (Figure 5F) to illustrate the shift in AR sensitivity observed upon RARγ loss.

Together, these data suggest that RARγ expression significantly regulates AR function in normal prostate cells independently of exogenous retinoid ligand. Specifically, RARγ loss impacts both phenotypic and transcriptomic androgen responses which manifest as a dampened response to DHT that normally slows proliferation and induces differentiation.

### Elevated miR-96 drives reduced RARγ expression and associates with aggressive prostate cancer

The frequent downregulation of *RARG* reflected neither mutation nor copy number variation^45^ and therefore we considered epigenetic mechanisms. In the first instance we assess DNA methylation data available in TCGA-PRAD cohort, but found no evidence for altered methylation at *RARG* TSS loci, although we found stronger evidence for increased CpG methylation associated with the paralogs RARα and notably RARβ (**Supplementary Figure 7**). Therefore, we considered miRNA that may target RARγ, and used *in silico* prediction tools^20^ to define a cohort of miRNAs that target the commonly downregulated NRs (e.g. *RARG, GR*) we previously identified in the TCGA-PRAD and MSKCC cohorts ^45^.

**Figure 7:**
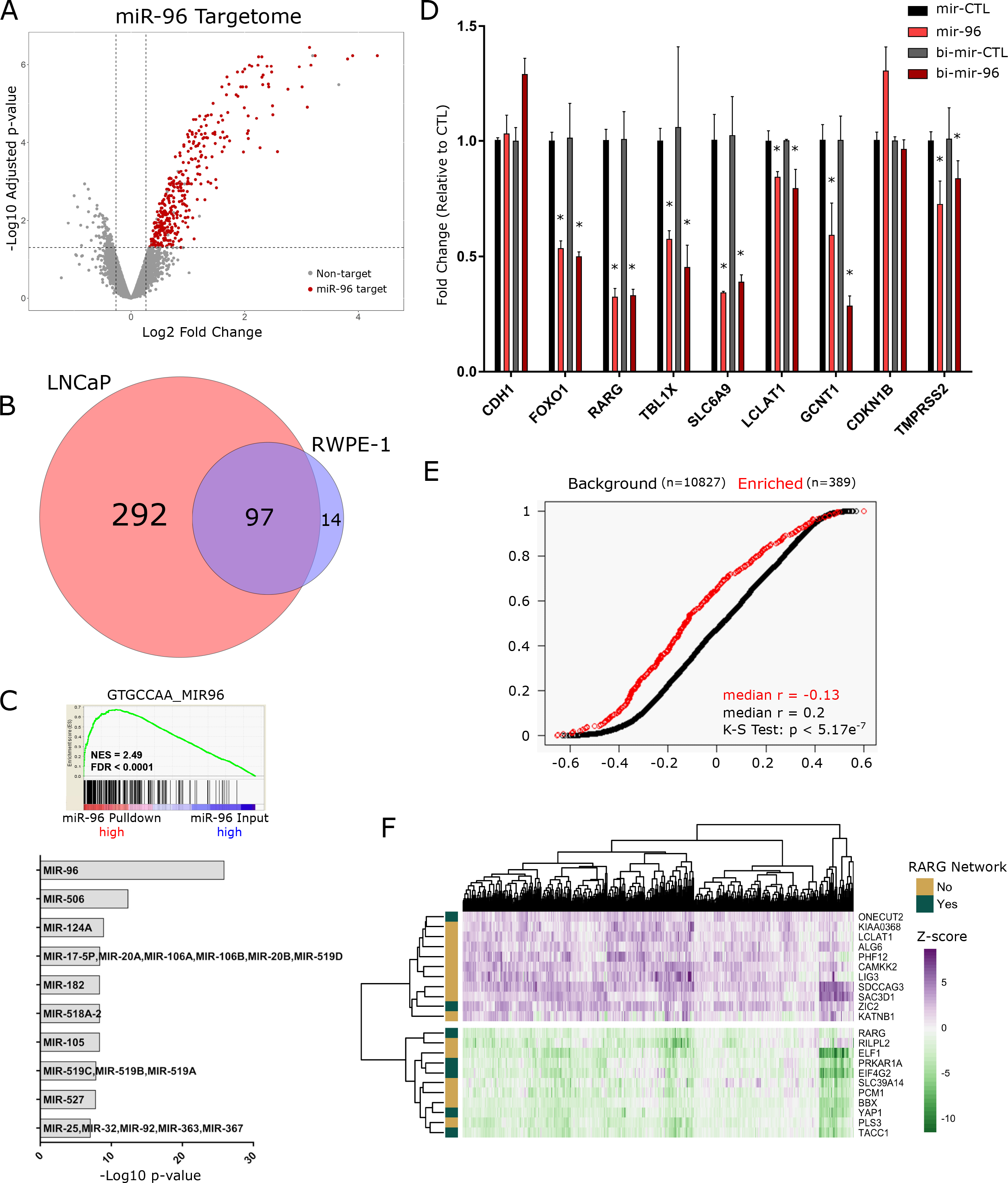
The miR-96 targetome centers on a RARγ-network. (**A**) LNCaP cells were transfected in triplicate with bi-miR-96 or bi-cel-miR-67 (non-targeting control, bi-miRCTL) (30 nM) for 24 hours prior to cell lysis. Input (5% of cell lysate) and streptavidin pulldown RNA were isolated and analyzed by Illumina microarray (Illumina HT12v4) (DESeq2). Volcano plot depicting the enrichment of all genes in bi-miR-96 pulldown over input in LNCaP cells. Genes marked in red (n = 389) were considered experimentally determined miR-96 direct targets, as they were significantly enriched (FC > 1.2, p.adj < 0.05) in bi-miR-96 pulldown but not in bi-miR-CTL pulldown. (**B**) Venn diagram representing the overlap of miR-96 targetomes in LNCaP and RWPE-1 cells. (**C**) The top significantly enriched GSEA pathway from unbiased enrichment analysis of bi-miR-96 samples in LNCaP cells (pulldown/input) (top), and summary of top miRNA seed sequence matches in 3’UTR regions of experimentally determined miR-96 targets using the GSEA-MSigDB microRNA targets tool (bottom). (**D**) Either bi-miR-96, or nonbiotinylated miR-96 mimics, or respective control mimics, were transfected (30nM) in LNCaP cells for 48 hours and target gene expression examined in triplicate. *CDH1* was assessed as a negative control, and specific targets were chosen either as they were either previously validated (*FOXO1*, *RARG*) and/or were significantly enriched in bi-miR-96 pulldown profiling. Significance of difference in target transcript level between biotinylated and non-biotinylated miR-96 relative to respective controls is noted (* < 0.05). (**E**) Cumulative distribution plot comparing the correlations (Pearson’s r) between miR-96 and all detectable genes from bi-miR-pulldown assay (n = 10,827, black) in TCGA-PRAD cohort samples compared to the correlations between miR-96 and identified miR-96 targets (n = 389, red) across the same samples. Significant difference between distributions is determined by Kolmogorov-Smirnov test. (**F**) Heatmap depicting expression of annotated miR-96 targetome genes in the TCGA-PRAD cohort (pheatmap). Genes were filtered to identify those genes that were altered by more than +/− 2 Z scores in 35% of tumors relative to normal samples. Functional relationship of identified miR-96 target genes to the previously established RARγ network are indicated.

Specifically, 61 putative NR-targeting miRNAs were identified (**Supplementary Figure 8**). We used a bootstrapping approach to test if these 61 miRNA were altered at the level of expression in PCa cohorts. Collectively, these 61 miRNA were more significantly elevated in PCa samples than predicted by chance (p.adj = 0.02) and were reciprocal to the reduction of NR mRNA expression^45^. The miR-96-182-183 cluster was amongst the most commonly upregulated miRNAs. Although similar, base substitutions in the targeting regions of each miRNA suggest unique functionality (**Supplementary Figure 9A**). Only miR-96 and miR-182 have predicted target sequences in the *RARG* 3’UTR (**Supplementary Figure 9B**).

**Figure 8:**
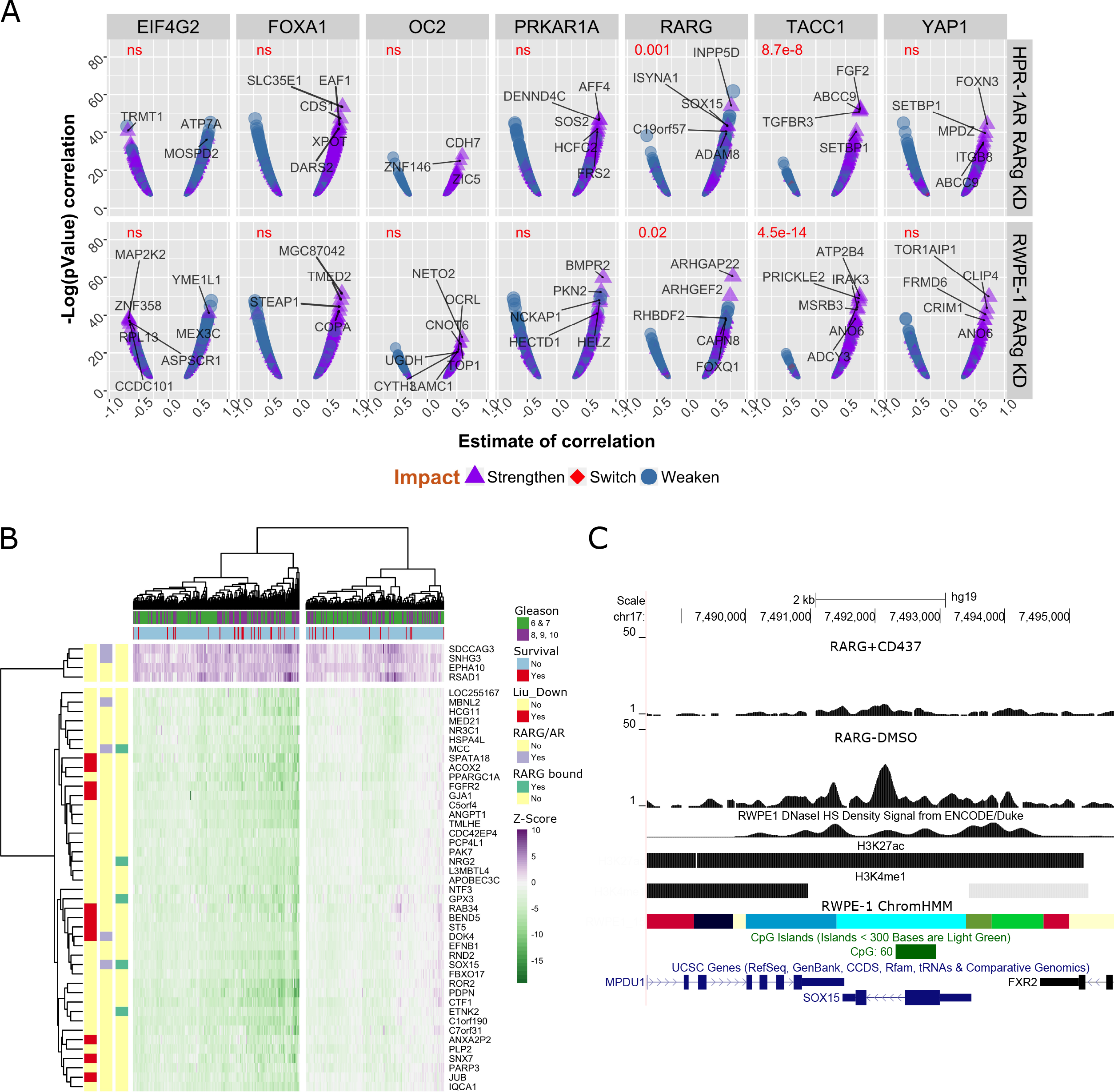
A miR-96/RARγ/TACC1 network associates with recurrent prostate cancer. (**A**) For each of the seven miR-96/RARγ network associated genes (*EIF4G2*, *ONECUT2*, *PRKAR1A*, *RARG*, *TACC1*, *YAP1*, *ZIC2*) plus *FOXA1* the Pearson r correlation was determined against all expressed genes within upper quartile of miR-96 expressing TCGA-PRAD tumors (miR-96_high_, n=123) and separately within the lower quartile (miR-96_low_, n=122). Significant correlations (q-value < 0.1) were selected for those genes that were RARg dependent in either RWPE-1 (Figure 3) or HPr1-AR (Figure 5). The shift in the strength of the correlation between miR-96_low_ compared to miR-96_high_ tumors was calculated, the correlations in miR-96_low_ tumors are visualized indicating if the correlation was strengthened, weakened, or switched direction between miR-96_low_ compared to miR-96_high_. A t-test (Welch Two Sample t-test) assessed the difference of the statistical strength of correlations between those that were positive and strengthened, compared to those correlations that were negative and weakened between miR-96_low_ and miR-96_high_ tumors. Genes that displayed the most significant change in correlation between miR-96_low_ and miR-96_high_ tumors are indicated in each panel. There were fewer than 50 significant correlations with *ZIC2* and is omitted. (**B**) TCGA-PRAD cohort tumor samples were separated based on expression of RARγ, TACC1 and miR-96 (based on lower/upper quartile expression) to generate RARγ/TACC1_low_, miR-96_high_ (n = 60) and RARγ/TACC1_high_, miR-96_low_ (n = 66) tumors and differential expression undertaken. Filtering the 1728 differentially expressed genes between these tumors by Z scores as described in Figure 7F revealed which genes were most altered in the TCGA-PRAD cohort. Unsupervised hierarchical clustering of gene expression identified a group of tumors that after adjusting for age significantly associated with worse disease free survival (hazard ratio 2.23, 95% CI 1.58 to 2.88, p = 0.015), and also clustered high Gleason score tumors (p = 0.012) (survival). Individual genes were annotated; RARG Bound are those identified by RARγ ChIP-Seq (Figure 4); RARG/AR are those displaying RARγ-dependent DHT regulation (Figure 5); and enrichment in the Liu prostate cancer gene set (Liu_Down). (**C**) Representative image of the *SOX15* locus, which is a RARγdependent AR regulated target, is bound by RARγ, and also shows a significantly stronger correlation with RARγ in miR-96_low_ tumors relative to miR-96_high_ tumors (**A**, upper panel). ChromHMM (cyan= TSS flanking; grass green = low transcription; bright green = Active enhancer; red = Transcriptional enhancer; blue = Active TSS; pale yellow = Poised transcription), CpG island, histone (GSE63094) and DNase sensitivity (GSM1541008) tracks are also shown.

In both the MSKCC and TCGA-PRAD cohorts, *RARG* significantly and negatively correlated with miR-96-182-183 cluster members, but most strongly with miR-96 (Figure 6A). Expression analyses in a panel of prostate cell lines (Figure 6B) revealed miR-96 was elevated and *RARG* reduced relative to non-malignant RWPE-1 and HPr1AR cells in all PCa cell lines examined. Profiling in PCa mouse models revealed reduced *Rarg* in both Pten^−/−^ model^40^ and to a greater extent in the TRAMP model^24^ but not in Hi-MYC mice^21^ relative to wild type mice (**Supplementary Figure 10A**). Indeed, *Rarg* expression decreased with PCa development in TRAMP mice relative to age-matched wild-type controls (C57BL/6), and inversely correlated most significantly with elevated miR-96 expression (Pearson’s r = −0.88) (Figure 6C, **Supplementary Figure 10B**). Furthermore in the TRAMP model, significant expression changes were observed as early as 10 weeks of age, when the mice display prostate epithelial hyperplasia. Finally, the expression changes of *RARG* and miR-96 cluster members were validated in an independent cohort of 36 matching tumor/normal prostate tissue pairs obtained from Roswell Park, corroborating observations in TCGA-PRAD and MSKCC cohorts, that furthermore associated with Gleason sum (**Supplementary Figure 11A-B**). Additionally, miR-96 expression in the TCGA-PRAD cohort associated with Gleason sum and clinical stage (**Supplementary Figure 11C**) and stratified patients based on shorter disease free survival (p = 0.018, **Supplementary Figure 11D**). Together, these findings in murine and human prostate tissues suggest that increased miR-96 expression occurs early and is sustained in PCa development, correlates strongly with loss of *RARG* and associates with aggressive PCa outcomes.

Two miR-96 recognition elements were identified in the *RARG* 3’UTR region (**Supplementary Figure 9B**). Ectopic overexpression of miR-96 resulted in significant loss of RARγ mRNA and protein in both RWPE-1 and LNCaP cells (Figure 6D, **Supplementary Figure 12A-C**). To demonstrate miR-96 interacts directly with RARG we developed and transfected a *RARG* 3’UTR containing luciferase vector into cells and measured a significant reduction in luciferase activity with the addition of exogenous miR-96 mimic (Figure 6E). Next we tested how miR-96 modulated the RARγ ligand-dependent induction and repression of target genes previously identified (Figure 6F). Indeed, 6 out of 7 CD437 induced (*CYP26A1*, *CYP26B1*, *FOXA1*, *PPARG*) or repressed (*FOXP1*, *KRT5*) genes were significantly dampened by the presence of miR-96, suggesting a reduction in RARγ sensitivity. Furthermore, we found that antagomiR-96 treatment in combination with miR-96 mimic rescued expression of *RARG*, and thus rescued RARγ regulation in almost all cases. Finally, we tested how miR-96 overexpression impacted ATRA regulation of a previously identified^67^ RARγ target gene, transglutaminase (*TGM4*). The induction of *TGM4* was dampened by both siRNA targeted to RARγ and by miR-96 mimic (**Supplementary Figure 13B**).

The impact of miR-96 overexpression on cell viability and cell cycle distribution were tested. Transfection with miR-96 mimics led to a significant increase in viability of LNCaP cells (p = 2.5e^−3^) (Figure 6G). These findings reflect phenotypes observed in RARγ knockdown cells, where reduced expression of the receptor associated with increased viability (Figure 3). Exogenous miR-96 also altered the cell-cycle distribution. In LNCaP cells there was also a loss of cells in G_1_/G_0_ phase (53.2% relative to 64.8%) and increase in S and G_2_/M phase (**Supplementary Figure 13A**). Cell cycle shifts in LNCaP cells transfected with miR-96 mimics did not mirror those observed in RARγ knockdown cells, and furthermore no significant changes were noted in RWPE-1 cells, suggesting that miR-96 related phenotypes also involve additional gene targets other than RARγ.

Together, these findings suggest elevated miR-96 is common in PCa and targets and suppresses *RARG* expression in order to dampen RARγ function, an interaction, which in mouse models at least, occurs early in PCa initiation. In LNCaP cells miR-96 mimics can increase viability and in PCa tumors miR-96 levels associate with higher Gleason score, clinical stage and predict disease recurrence.

### MiR-96 targets a network of RARγ interacting co-factors

While these data support miR-96 targeting of RARγ, other studies have identified additional targets in prostate and other cancers including the pioneer factor FOXO1^5, 28, 56, 87^. Therefore, to identify all miR-96 targets (the miR-96 targetome) in an unbiased manner, we undertook a biotin-miRNA (bi-miR) capture and streptavidin pull down approach coupled with microarray analyses^75^. Biotin labelling of miR-96 mimics did not interfere with either transfection or knockdown efficacy of target gene expression (**Supplementary Figure 14A-B**), and was able to capture known (e.g. *RARG* and *FOXO1*) and predicted (*TBL1X*) miR-96 targets (**Supplementary Figure 14C**).

In the first instance, processing of the bi-miR-96 and control mRNA libraries and PCA analyses revealed strong separation of experimental groups (**Supplementary Figure 14D**). Subsequently, comparing bi-miR-96 pulldown to input RNA (FC > 1.2, p.adj < 0.05) and to non-targeting control (bi-miR-CTL) pulldown revealed 111 and 389 miR-96 targets in RWPE-1 and LNCaP cells, respectively, which were largely shared, but also had unique genes (Figure 7A-B, **Supplementary Figure 14E**). Amongst these targets,3’UTR miR-96 target site motifs (GTGCCAA) was the most highly enriched miRNA motif (Figure 7C) (GSEA in LNCaP p.adj = 1.24e10^−26^, RWPE-1 p.adj = 1.70e10^−9^), underscoring the efficacy of this method to capture miR-96 targets. RT-qPCR validation of bi-miR-96 capture methodology in LNCaP cells independently confirmed the capture of known and novel targets identified from profiling efforts (**Supplementary Figure 15F**).

The capacity of miR-96 to suppress target gene expression was also assessed by independent transfections of biotinylated and non-biotinylated miR-96 mimics. In LNCaP cells, 6 out of 7 identified miR-96 targets were significantly downregulated 48 hours post miR-96 or bi-miR-96 transfection, relative to respective non-targeting controls, while a negative control target *CDH1* was not affected (Figure 7D). Intriguingly, the single transcript not downregulated in this experiment, p27 (encoded by *CDKN1B*), showed significantly reduced protein level upon miR-96 overexpression, suggesting that in some cases the result of miR-96 binding is translational inhibition not degradation of transcript (**Supplementary Figure 15G**). Similar regulation of targets was observed in RWPE-1 cells (**Supplementary Figure 15H**).

Furthermore, in LNCaP cells *FOXO1* was neither significantly enriched in the bi-miR-96 fraction upon microarray analysis and was only validated by RT-qPCR in RWPE-1 cells. Also surprisingly, *RARG* transcript was not significantly detectable via microarray in either LNCaP or RWPE-1 cells, but did validate by RT-qPCR in both cell types. It was revealed that the Illumina HT12v4 Bead Chip array contains only a single exon probe targeting a single *RARG* isoform. *RARG* expression in LNCaP cells has been reported suggesting that the microarray finding is a false negative one.

Functional annotation of the miR-96 targetome in LNCaP cells, based on GO terms, revealed roles for miR-96 in governing cell cycle progression, metabolism, cell morphology and microtubule organization (**Supplementary Figure 15A**). These observations reflected the impact of miR-96 on cell cycle and included *CDKN1B*, *CDK2*, *DICER1* and transforming acidic coiled-coil protein 1 (*TACC1*). Validation of the regulation of selected cell-cycle regulators and transcription factors (e.g. *BTBD3*) was undertaken in LNCaP cells after transfection with miR-96 mimics. Expression of several of these (e.g. *TACC1* and *BTBD3*) were suppressed (**Supplementary Figure 15B,C**).

To assess whether experimentally determined targets had expected relationships with miR-96 in clinical samples, the TCGA-PRAD data were examined again. In the first instance, we examined the correlation of expression between miR-96 and identified miR-96 targets. For example, there was a significant inverse correlation between miR-96 and *TACC1* in the TCGA-PRAD cohort (r = −0.54) (**Supplementary Figure 15D**). Comparing the distribution of these correlations of derived miR-96 targets against that of the background transcriptome revealed a significant shift toward negative correlations between miR-96 and its targets, supporting a functional relationship (p = 5.17e^−7^) (Figure 7E).

Finally, we measured the distorted expression of the miR-96 targetome in the TCGAPRAD cohort. Stringent filtering of expression of the miR-96 targetome genes (+/−2 Z-scores in 35% of tumors relative to normal prostate tissue) identified 22 genes with significantly altered expression (Figure 7F). Seven of these 22 genes included *RARG* itself as well as six other genes that are either established (*TACC1*^27^, *ZIC2*^50^, *PRKAR1A*^65^, *EIF4G2*^62^) or putative (*ONECUT2*^35^ and *YAP1*^29^) RAR co-regulators. Interestingly, the motif for ONECUT2 was enriched in the experimentally derived RARγ cistrome and has been previously identified as a miR-96 target^92^.

### The miR-96 governed RARγ network associates with recurrent prostate cancer

Several lines of evidence support the concept that miR-96/RARγ is a significant signaling axis in the prostate; in the absence of exogenous retinoid ligand, experimentally manipulating expression levels of RARγ controls gene expression patterns and governs the sensitivity of the AR to DHT stimulation; miR-96 modulates the expression levels of RARγ and a number of interacting co-factors as well as RARγ associated regulatory functions; the expression of miR-96 and RARγ are reciprocal in murine and human tumors; both the gene targets of RARγ (e.g. *CYP11A1*) and miR-96 itself predict prostate cancer progression in patients.

To dissect the impact of the miR-96 regulation of RARγ further we examined how the correlations of each of the seven RARγ network genes were altered by expression levels of miR-96. Specifically, we identified the significant correlations between each of the seven RARγ network genes (*EIF4G2*, *ONECUT2*, *PRKAR1A*, *RARG*, *TACC1*, *YAP1*, *ZIC2*^50^) alongside *FOXA1* with all RARγ dependent genes in the upper (miR-96_high_, n=123) and lower (miR-96_low_, n=122) quartile miR-96 expression TCGA-PRAD tumors. From these distributions we next calculated how the strength of each correlation changed between miR-96high and miR-96_low_ tumors. That is, we reasoned that correlations between RARγ network and RARγ target genes would significantly strengthen as miR-96 levels decreased. Testing these relationships revealed that this was true only for RARG and TACC1 (Figure 8A). For example the difference in the strength of the correlation between TACC1 and the genes altered by RARγ knockdown in HPr1-AR or RWPE-1 was measured between patients within the lower and upper quartile of miR-96 levels (Welch two sample t-test; p.adj = 8.7e^−8^, and 4.5e^−14^ with HPr1-AR or RWPE-1 knockdown genes, respectively).

Therefore we sought to establish the combined significance of miR-96/RARγ/TACC1 axis in PCa by further segregating tumors in the TCGA-PRAD cohort into low quartile expression of RARγ and TACC1 (RARγ/TACC1_low_), and miR-96_high_ expression. These were compared to the reciprocal (RARγ/TACC1_high_, miR-96_low_) (Figure 1) for DEG analysis. This approach identified 1728 genes, which were then overlapped with the gene sets developed throughout this study to establish how they related to the various conditional RARγ networks identified. Using hypergeometric testing, we measured the significance of overlaps between the *in vitro* derived gene sets and those identified in the miR-96/RARγ/TACC1 axis in TCGA-PRAD. For example, 118 genes were shared between RARγ knockdown in RWPE-1 and HPr1-AR overlapped (p = 7.6e^−12^), and 119 genes were shared between miR-96/RARγ/TACC1 axis genes RARγ-dependent genes in HPr1-AR (p = 5.3e^−12^) (**Supplementary Figure 16A**). These findings suggest that the underlying biology of RARγ dissected in cell line models was detectable in human tumors.

We reasoned that if the genes identified as dependent upon the miR-96/RARγ/TACC1 axis were critical in PCa, they would be commonly distorted and associate with aggressive disease. Filtering the ~1700 differentially expressed genes in RARγ/TACC1_low_, miR-96_high_ tumors (altered by +/− 2 Z scores in more than 45% of tumors relative to normal prostate tissue) revealed 47 genes. Unsupervised hierarchical clustering of the expression of these genes identified a group of patients in the TCGAPRAD cohort that after adjusting for age significantly associated with high Gleason score tumors, (p = 0.012) and worse disease free survival (hazard ratio 2.23, 95% CI 1.58 to 2.88, p=0.015) (Figure 8B).

Genes were also annotated for whether they were identified as a RARγ cistrome gene (RARG bound); DHT-regulated RARγ-dependent genes (RARG/AR); and member of the highest rank GSEA term (LIU_PROSTATE_CANCER_DN; a cohort of genes significantly down-regulated in PCa^43^). Five RARγ cistrome genes were revealed, all down-regulated, including the tumor suppressor *MCC*^38^ and other regulators of cell-fate such as *GPX3* and *SOX15*^78^ and *MCC* and *SOX15* were also DHT-regulated RARγ-dependent genes (Figure 8C). Furthermore the correlation of between *RARG* and *SOX15* was one of the top most strengthened correlations (Figure 8A, **upper panel**). These findings support the concept that the miR-96/RARγ/TACC1 axis can be detected in PCa cohorts, and contains genes that associated with AR cross talk and the risk of disease recurrence.

## DISCUSSION

The rationale for the current study arose from our prior observation of the common down-regulation, but neither the deletion nor mutation, of RARγ in PCa^45^. Therefore we modulated RARγ expression in three different prostate models to examine the consequences on cell viability and transcriptional responses. These data were complemented by RARγ ChIP-Seq data, and publicly available cistromic data sets to reveal that RARγ was significantly bound at gene enhancers and could significantly regulate the sensitivity of AR signaling. We next identified miR-96 targeting as a significant cause of down-regulation of RARγ and further revealed the miR-96 targetome to include a number of RARγ interacting co-factors. Finally, we revealed that tumors with up-regulated miR-96 and down-regulated RARγ were significantly more likely to recur following initial therapy.

Developmental roles for RARγ have been identified in skin, skeletal and reproductive systems, including the prostate^44^. Indeed, *Rarg−/−* mice display prostate epithelial squamous metaplasia due to improper glandular formation^44^. More broadly, RARγ has emerged as an important transcription factor. For example, the Roadmap Epigenome consortium identified RARγ as a member of the most highly enriched in transcription factor-enhancer interactions across the human epigenome^68^ and others have considered a role for the receptor to govern pluripotency^70^. *RARG* has also been suggested to be a tumor suppressor in keratinocytes^15^.

We have now identified that the actions of RARγ are substantial in the absence of exogenous retinoid ligand, and potentially more biologically significant than the impact of retinoid exposure. Similar findings are also found in *Rarg*^−/−^ mice. Specifically, we analyzed publicly available microarray studies undertaken in *Rarg*^−/−^ murine ES cells and also F9 cells^36^. This analysis, in a genetically clean system, revealed similar patterns resulting from depletion of RARγ namely deletion of receptor had the largest impact on gene expression, compared to treating wild-type cells with exogenous retinoid (**Supplementary Figure 16B**).

There is a significant pre-clinical literature dissecting RARs in solid tumors including PCa^10, 33^ with the goal of developing so-called differentiation therapies^52^. The goal of these approaches was to exploit retinoid signaling in PCa as a therapeutic approach to temper and subdue AR signaling. The frequently observed CpG methylation of the RARβ perhaps underscores the significance of these receptors in PCa progression^71^. However, the Phase I/II clinical trials of retinoids alone^53^ or in combination^64^ in patients with advanced PCa met with only limited clinical success^23, 82^. Furthermore there are ambiguous relationships between serum levels of retinoid metabolites and PCa^6, 59, 81^. Nonetheless, there appears to be a significant biological relationship between RARγ and PCa^67^.

Given that expression of RARγ regulated genes such as *SOX15* associated with worse disease outcome, we propose that the reduction in RARγ is clinically significant because it impacts the RARγ gene-regulation capacity in the absence of exogenous retinoid. These findings are also perhaps compatible with other receptors such as RARβ being more prominent in the sensing of exogenous ligand in PCa cells^10^.

A key finding from the current study is the role for RARγ regulation of AR signaling in PCa. Others have implicated RARs generally with PCa genomic states^60^, or dissected RARγ cross-talk with AR signaling at the candidate gene level^67^. The current study therefore extends these earlier findings to develop a genomic understanding of the cellular and clinical significance of the role of RARγ in the absence of exogenous retinoid ligand. These findings also reflect an emerging understanding of the RARγ functional crosstalk with steroid receptor signaling in breast cancer^30^.

Genomic RARγ binding in RWPE-1 cells, in the absence of exogenous ligand, significantly overlapped with histone modifications that defined active enhancers and DNase sensitivity (also in RWPE-1) and AR binding and p65 binding (in LNCaP). However RARγ binding did not overlap with VDR binding in LNCaP. TCGA investigators^12, 14^ and others^25^ have identified common translocations between *TMPRRS2* and *ETS* oncogenes in PCa^69, 79^. *TMPRSS2* is androgen responsive, and its translocation leads to androgen regulation of *ETS* oncogenes, which in turn act as cancer-drivers^84, 88^. VCaP is a TMPRSS2 fusion positive cell therefore we examined how RARγ overlapped with AR and the ETS family member, ERG, in these cells. Interestingly, whilst RARγ overlapped with AR in LNCaP cells, it did not overlap with either AR or ERG in VCaP cells, and may reflect the impact of ERG on the genome.

RARγ loss profoundly altered AR regulated events including DHT-mediated antiproliferative effects^16^ and reduced the sensitivity of the DHT-dependent transcriptome, notably by restricting the capacity of AR to repress MYC signaling. Recently MYC has been shown to antagonize AR function in prostate cells^32^, and the current study suggest that RARγ regulates these events, in part through direct binding at the MYC locus.

It is possible that the changing expression levels of RARγ impacts the expression of cofactors that can be shared with the AR. To address this possibility we generated a comprehensive list of ~1900 transcription factor co-regulators by mining FANTOM, Uniprot and the HGNO data bases and tested how these overlapped with the gene expression following RARγ knockdown in each of RWPE-1, HPr1-AR and LNCaP. (**Supplementary Figure 17**). This approach identified co-activators appeared to be commonly enriched in gene sets that resulted from changes to RARγ (e.g. knockdown of RARγ in RWPE-1, n = 47), and DEGs associated with the miR-96/RARγ/TACC1 axis in TCGA PRAD tumors (n=49). This leads to the intriguing suggestion that RARγ levels regulates expression of a number of co-regulators, which in turn impact how the AR signals. These co-regulators included *FHL2*^55^, a co-regulator of FOXO1; *LPIN1*^8^ and *PPARGC1B*^1^, which interact with PPARs signaling; several negative co-regulators of WNT (*DKK1*, *DKK3*, *PRICKLE1*, *SFRP1*)^58^ and HIPPO (*WWTR1*)^42^ signaling. However in the context of the current study, the number of genes prohibits obvious intervention approaches (e.g. over-expression or knockdown).

Although miR-34c targets *RARG* in embryonic stem cells^93^, we found no significant relationship between miR-34c and RARγ in the current study. Rather we identified miR-96, which has been investigated in various cancer settings including in PCa, but not associated with RARγ^28, 83^. Furthermore, miR-96 is one of the 60 miRNAs whose expression most differentiated PCa clinical groups in TCGA-PRAD cohort^13^. The current study extends these reports of oncogenic actions of miR-96 to identify the RARγ network as a major biological and clinically-relevant target.

The inverse relationships between miR-96 and RARγ were robust and significant across cell models, murine PCa models and clinical cohorts, and equal or greater than with previously identified targets including *FOXO1*. Biochemical tests established that miR-96 binds directly to multiple target sites contained within the 3’UTR of *RARG* and using the bi-miR capture approach revealed ~400 direct targets enriched for predicted miR-96 target sequences, and identified targets were directly bound and downregulated upon miR-96 overexpression.

Experimental over-expression of miR-96 positively regulated proliferation and cell cycle in PCa cells reflecting previous reports^28^, but did not affect proliferation in non-malignant prostate cells. It is thus interesting to speculate that miR-96 acquires additional targets and subsequent functions within the malignant transcriptome, which coincides with the observation of 3-fold additional targets in malignant LNCaP cells relative to non-malignant RWPE-1, and impacts the control of cell cycle progression. Given that RARγ appears to function as an important AR co-regulator to promote differentiation, then its downregulation by miR-96 targeting suggests an oncogenic action that shifts the AR signaling capacity in localized PCa. Notably, miR-96 has previously been reported as androgen regulated in prostate cells^57^. Filtering miR-96 target genes to those most altered in PCa revealed that nearly one third (7/22) were within an RARγ-centric network of co-regulators, suggesting its regulation as an important aspect of miR-96 function.

Differential expression analyses in TCGA-PRAD cohort between tumors showing reciprocal expression of miR-96 and RARγ/TACC1 clustered aggressive tumors and predicted worse disease free patient survival (Figure 8B). That is, within the TCGA-PRAD cohort, 390 out of 498 tumors have data on disease free survival and Gleason score, and of these, 37 have recurrent disease. The differential expression of these genes determined by RARγ/TACC1 and miR-96 expression significantly predicts the risk of developing recurrent disease. We suggest therefore that this gene panel may be of interest to test in larger clinical cohorts or as part of meta-analyses.

Interestingly, recently TACC1 and RARγ interactions have been identified to be critical for neural development^89^. Several of the shared target genes identified in the current study are either under-explored in PCa (e.g. *SOX15* and *MCC*) or have only been investigated in metastatic PCa^80^ (*PPARGC1A*). Together these data strongly support the concept that the miR-96/RARγ is a significant and yet under-explored regulator of the AR and drives PCa progression.

Finally, we believe these studies may form a platform for a number of highly translational follow on studies. That this AR-centered transcriptional circuit is regulated by miR-96 adds diagnostic and therapeutic potential. In the first instance miR-96 may have significant diagnostic potential when measured in the tumor or serum of new PCa patients^2, 54, 72^, and have potential to be targeted with antagomir based therapies^34, 63^.

We also believe it is timely to revisit the previous clinical experience with targeting retinoid signaling in PCa. We propose that a lack of success in clinical exploitation of retinoid-based therapies in PCa reflects an under-appreciation of the role of RARγ to function in the absence of exogenous ligand. These functions are centered on the regulation of gene networks that relate to the regulation of AR function and broader impact on cell proliferation and differentiation. It is reasonable to propose that there may also exist under-appreciated genomic functions for other Type II nuclear receptors (e.g. VDR, PPARs, FXR, LXRs, CAR)^11, 31, 86^ that exert genomic functions in the absence of exogenous ligand and are potential targets in cancer.

## MATERIALS AND METHODS

### Data analyses, integration and code availability

All analyses were undertaken using the R platform for statistical computing (version 3.1.0) and a range of library packages were implemented in Bioconductor^26^. All code is available upon request from the corresponding author.

### Cell culture and materials

All cells were maintained at 37°C and 5.0% CO_2_ using a cell culture incubator with UV contamination control (Sanyo). RWPE-1, RWPE-2, PNT2, and HPr1AR cells were maintained in Keratinocyte-Serum Free Medium (KSF) containing human recombinant Epidermal Growth Factor 1-53 (EGF 1-53) and Bovine Pituitary Extract (BPE). LNCaP, LAPC4, EAA006, MDAPCa2b, LNCaP-C42, 22Rv1, PC3, and DU145 cells were maintained in RPMI 1640 Medium containing 10% FBS. All media was supplemented with 100 U/mL Penicillin-Streptomycin. ATRA, CD437 and DHT (sigma) were kept in DMSO or EtOH stocks, and diluted to 1000x stocks in DMSO prior to treatments. All cells were sourced from ATCC.

### Stable knockdown of RARγ

Knockdown of RARγ in RWPE-1, LNCaP and HPr1AR cells was achieved by stable selection after transduction with lentiviral shRNA constructs targeting *RARG*. Two targeting constructs (V2LHS_239272, V2LHS_239268) and one non-silencing control construct were selected from the V2LHS pGIPZ based lentiviral shRNA library (Thermo Fisher Scientific) for testing. Viral packaging and cellular infection was performed through the RPCCC shRNA Resource. All pGIPZ containing cells were subsequently maintained in media supplemented with puromycin (2µg/mL), including during all experiments.

### BAC-RARG-EGFP

BAC-RARG-EGFP construct (CTD-2644H7) was a generous gift of Dr. Kevin White (University of Chicago). RWPE-1 cells were transfected with BAC-RARG-EGFP construct using Lipofectamine^®^ 3000 and selected with G418 and consistently maintained under antibiotic selection for all subsequent passaging of cells and also during experiments.

### Cell viability

Bioluminescent detection of cellular ATP as a measure of cell viability was undertaken using ViaLight^®^ Plus Kit (Lonza Inc.) reagents. Cells were plated at optimal seeding density to ensure exponential growth (4×10^3^ cells per well) in 96-well, white-walled plates. Wells were dosed with agents to a final volume of 100 µl. Dosing occurred at the beginning of the experiment, and cells were incubated for up to 120 hours. Luminescence was detected with Synergy^TM^ 2 multi-mode microplate reader (BioTek^®^ Instruments). Each experiment was performed in at least triplicate wells in triplicate experiments.

### Cell cycle analysis

Cells were seeded in 100 × 20mm sterile, polystyrene tissue culture dishes (Corning Inc.) at densities previously optimized for each cell line, and were allowed exponential growth with our without ligand for specified times (RWPE-1: 1µM ATRA or 10nM CD437 for 72 hours with re-dosing after 48 hours, LNCaP: 1µM ATRA or 250nM CD437 for 120 hours with re-dosing after 48 and 96 hours). Mid-exponential growth cell cultures were harvested with accutase (Invitrogen), fixed and stained with propidium iodide buffer (10 µg/ml propidium iodide, 1% (wt/vol) trisodium citrate, 0.1% (vol/vol) TritonX-100, 100 µM sodium chloride) for 1 hour, on ice, in the dark. Cell-cycle distribution was determined utilizing FACSCalibur^TM^ Flow Cytometer (Becton-Dickinson) and analyzed with ModFIT 3.1 SP3 cell-cycle analysis software.

### RT-qPCR

Quantitative real-time reverse transcription–polymerase chain reaction (RT-qPCR) was employed for detection of target mRNA transcripts. Total RNA was isolated via TRIzol^®^ reagent (Thermo Fisher Scientific) for candidate detection or by use of the High Pure RNA Isolation Kit (Roche), following manufacturer’s protocols. Complementary DNA (cDNA) was prepared using iScript^TM^ cDNA Synthesis Kit (Bio-Rad) according to manufacturer’s protocol. Relative gene expression was subsequently quantified via Applied Biosystems 7300 Real-Time PCR System (Applied Biosystems), for both TaqMan and SYBR Green (Thermo Fisher Scientific) applications. All targets were detected using either pre-designed TaqMan Gene Expression Assays (Thermo Fisher Scientific; *ALDH1A2, AR, CDH1, CDKN1B, CRABP1, FOXO1, GAPDH, GCNT1, Gusb, KRT8, LCLAT1, PPARG*, *RARA, RARB, RARG*, *Rarg, SLC6A9, TBL1X, TGM4, TP63*), pre-designed PrimeTime qPCR primers (IDT; *BTBD3, CYP26A1, EIF4G2, ELF1, KRT10, KRT18, LRAT, MORF4L1, PAFAH1B1, PRKAR1A, SMAD4, TACC1, USP9X, ZNF430*) or custom designed qPCR primers (IDT) using a final primer concentration of 500nM. All primers for use with SYBR Green application were tested for specificity by melting curve analysis with subsequent product visualization on agarose gel. All RT-qPCR experiments were performed in biological triplicates, with at least technical duplicates. Fold changes were determined using the 2^−ΔΔCt^ method, where ΔCt was calculated as the Ct of the target gene minus the Ct of endogenous control, and ΔΔCt was calculated as the difference between experimental group and respective control group. Significance of experimental comparisons was performed using Student’s t-test.

### Immunoblotting

Total cellular protein was isolated from exponentially growing cells for determination of target protein expression. Cells were harvested, then washed in ice cold PBS before lysing in ice cold RIPA buffer (50mM Tris-HCl pH 7.4, 150mM NaCl, 1% v/v Triton X-100, 1mM EDTA pH 8.0, 0.5% w/v sodium deoxychlorate, 0.1% w/v SDS) containing 1x cOmplete Mini Protease Inhibitor Tablets (Roche). Protein concentrations were quantified using the DC Protein Assay (Bio-Rad) as per manufacturer’s instructions. Equal amounts of proteins (30-60µg) were separated via SDS polyacrylamide gel electrophoresis (SDS-PAGE) using precast 10% polyacrylamide gels (Mini-Protean TGX, Bio-Rad). Proteins were transferred onto polyvinylidene fluoride (PVDF) membrane (Roche) for 80V for 1.5 hours. Post transfer, membranes were blocked with 5% non-fat dry milk (NFDM) for 1 hour at room temperature. Blocked membranes were probed with primary antibody against RARγ (sc550, Santa Cruz), GFP (ab290, Abcam), p27 (D37H1, Cell Signaling), or GAPDH (2118s, Cell Signaling) either overnight at 4°C or for 3 hours at room temperature. Primary antibody was detected after probing for 1 hour with HRP-linked rabbit anti-mouse IgG (P0161, Dako) or goat anti-rabbit IgG (P0448, Dako) secondary antibody at room temperature using ECL Western Blotting substrate (Pierce). Signal and quantification was performed using the ChemiDoc XRS+ system (Bio-Rad).

### Expression determination in PCa mouse models

Snap frozen tissue and/or RNA from previously harvested normal or malignant prostate tissues of Hi-MYC, PTEN−/− and TRAMP (5 tumors at each of 6, 8, 10, 15, 20-25 week) models, as well as from age-matched wild type (10 prostates from FVB:BL6, C57BL/6) mice, was obtained from the lab of Dr. Barbara Foster at RPCCC. Relative expression of *Rarg* and microRNAs was determined in these tissues by RT-qPCR after normalization to *Gusb* or RNU6B, respectively.

### miRNA mimic, antagomiR and siRNA transfection

Ectopic overexpression of miRNA was achieved by transient transfection of mirVana^®^ miRNA mimics (30nM) (Ambion). Inhibition of miRNA was achieved by transient transfection of Anti-miR^TM^ miRNA Inhibitor (30nM) (Ambion). For transient silencing of targets, siRNA was employed using Silencer^®^ predesigned siRNAs (30nM) (Ambion). Transfection was accomplished by use of Lipofectamine^®^ 2000 or Lipofectamine^®^ 3000 by manufacturer’s instructions. Concentrations of miRNA mimics and transfection reagents were optimized using BLOCK-iT™ Alexa Fluor^®^ Red Fluorescent Control (Ambion) as well as by subsequent miRNA specific RT-qPCR (TaqMan MicroRNA Reverse Transcription Kit, Thermo Fisher Scientific) and RT-qPCR of target mRNAs to assess efficiency of transfections.

### RPCCC and TCGA PCa samples

RNA was obtained from a cohort of 36 men who underwent radical prostatectomy at RPCCC and de-identified tissue made available through the Pathology Resource Network (PRN) and Data Bank and BioRepository (DBBR)^3^ shared resources at RPCCC. Areas of histologically normal tissue and areas with high percentages of neoplastic tissue were isolated and RNA extracted. De-identified patient codes were matched with Gleason scores for each sample, were made available through the RPCCC Clinical Data Network (CDN).

PCa data sets were downloaded through cBioPortal. The MSKCC data set (n=109) was in a tumor-normal Z score format and the TCGA-PRAD cohort (n=498; 390 with progression data) was also converted to this format. Expression of overlapping genes were filtered (`genefilter`) as indicated before visualization and clustering of expression (pheatmap) and testing the relationship between expression and patient outcomes (`survival`).

### Molecular Cloning / Luciferase Assay

Luciferase assay was employed to assess direct targeting of miR-96 to the full length (FL) RARG 3’UTR or individual predicted target sites (t1, t2) within the RARG 3’UTR. RSV-pGL3-basic firefly luciferase expressing vector (Addgene) was employed for this purpose, utilizing FseI and XbaI restriction sites located between the *luc+* gene and the SV40 pA tail. Primer sets were designed for PCR amplification of either the ~1100bp full-length (FL) *RARG* 3’UTR or two smaller regions (~300bp) within the *RARG* 3’UTR containing individual predicted miR-96 target sites (t1, t2). PCR amplicons were digested and ligated to RSV-pGL3-basic vector via T4 DNA Ligase in 1X T4 DNA Ligase Buffer (New England BioLabs, Inc.), using an insert to vector ratio of 2.6. Individual transformed *E.coli* colonies were expanded prior to plasmid isolation via E.Z.N.A^®^ Plasmid Mini Kit (Omega), and inserts verified by sanger sequencing. RWPE-1 cells were seeded in 96-well plates and then co-transfected with combinations of miR-96 or miR-control mimics (30nM), indicated RSV-pGL3 constructs (250ng), and pRL-CMV Renilla luciferase expressing vectors (25ng) via Lipofectamine^®^ 2000 for 48 hr. Media was subsequently removed, cells washed with PBS, and both firefly and Renilla luciferase activity measured by Dual-Glo Luciferase Assay System (Promega). For each construct (FL, t1, t2, empty), detected firefly luminescence was first normalized to Renilla luminescence, and then normalized luminescence was compared between miR-96 and miR-control transfected cells. Experiments were performed in technical triplicates, and experiment replicated a total of 5 times.

### Chromatin Immunoprecipitation

ChIP was performed in BAC-RARG-EGFP containing RWPE-1 cells in the presence of CD437 (10nM, 2hr) or DMSO as previously described^77^. Briefly, triplicate cultures of approximately 20x10^6^ cells were crosslinked with 1% formaldehyde solution, quenched with glycine (0.125 M) and harvested in cold PBS. Sonication of crosslinked chromatin was performed using a Bioruptor^®^ UCD-200 TM Sonicator (Diagenode) for a total of 15 minutes. Immunoprecipitation of sonicated material was performed with antibodies against eGFP (ab290, Abcam) or IgG (sc-2027x, santa cruz) for 16 hours, and antibody/bead complexes isolated with Magna ChIP^TM^ Protein A+G magnetic beads (Millipore). Complexes were washed, reverse crosslinked, and treated sequentially with RNase and proteinase K prior to DNA isolation. Sequencing (100bp single end, >30×10^6^average reads/sample) was performed at the RPCCC Genomics Shared Resource core facility. The RARγ cistrome was analyzed with `Rsubread/csaw`^49^, along with TF motif analyses (`MotifDb`). Binding site overlaps between RARγ and the other cistromes tested (`ChIPpeakAnno`^94^ and `bedtools`). Peak density plots were performed using the annotatePeaks.pl tool available from the HOMER (Hypergeometric Optimization of Motif EnRichment) suite.

### Biotin-miRNA pulldown

Approximately 20×10^6^ were transfected with either hsa-miR-96-5p or cel-miR-67 miRIDIAN miRNA mimics (30nM, 24hr) modified with a 3’biotin attached to the guide strand (Thermo Fisher Scientific) using Lipofectamine 3000 Transfection Reagent (Invitrogen). Harvested cell pellets were resuspended in cell lysis buffer (10mM KCl, 1.5mM MgCl2, 10mM Tris-Cl pH 7.5, 5mM DTT, 0.5% IGEPAL CA-630) containing SUPERase·In (Ambion) and 1x cOmplete Mini protease inhibitor (Roche) and cleared by centrifugation. 5% of cell lysate was collected to serve as lysate RNA input. Dynabeads MyOne^TM^ Streptavidin C1 (Thermo Fisher Scientific) were washed 3 times with bead wash buffer (5mM Tris-Cl pH 7.5, 0.5mM EDTA, 1M NaCl), and then blocked (1µg/µL bovine serum albumin, 1µg/µL Yeast tRNA, 50U/mL RNaseOUT) for 2 hr. Resuspended beads were added 1:1 to cell lysate, and mixed for 30 minutes. Bead-lysate mixtures were collected with a magnetic rack, and bead-bi-miR complexes washed a total 3 times with wash buffer. Bead-bi-miR complexes and input control samples were resuspended in water and purified using the Qiagen RNeasy^®^ Mini kit (Qiagen) according to manufacturer’s RNA clean-up protocol. To concentrate samples for downstream analyses, eluted RNA was brought up to a total volume of 500µL in H_2_O and filtered through Amicon Ultra-0.5mL Centrifugal Filters (EMD Millipore) according to manufacturer’s instructions. Subsequent amplification and labeling of 50ng of pulldown and input RNA was performed at the RPCCC Genomics Core Facility, using the Illumina^®^ TotalPrep RNA Amplification kit, including a 14 hour incubation for the IVT step. Hybridization of cRNA samples onto Illumina Human HT-12v4 bead arrays, and successive scanning and raw intensity measurement extraction were also performed at the RPCCC Genomics Core Facility.

### miRNA analyses

To reveal putative NR-targeting miRNA, miRWalk, a comprehensive database on predicted and validate miRNA targets, was employed. If at least 5 out of 9 algorithms positively predicted an interaction, it was considered in subsequent analyses. MiRNA expression was queried in PCa tissue samples and matched normal tissue from TCGA cohort data as previously described^45^. To examine if NR-targeting miRNA expression alterations significantly deviated from what would be expected by chance, bootstrapping approaches were utilized as previously described^45^.

### Microarray / RNA-seq analyses

Global changes in mRNA, biological triplicate samples per experimental condition were analyzed using Illumina microarray (Illumina HT12v4) or by RNA-seq (`limma`^66^ or `DESeq2`^47^). For RNA-seq data, raw sequence reads (75bp paired end, >45×10^6^ average reads/sample) were aligned to the human genome (hg19) using `tophat2`, and aligned reads translated to expression counts via `featurecounts`, followed by a standard `DESeq2` pipeline.

### Functional annotation gene sets

Functional annotations were performed using GSEA v3.0 and gene sets from the Molecular signatures database (MSigDB). Specifically, gene sets were compiled to assess enrichment of all BROAD Hallmark pathways, curated pathways (KEGG, BioCarta, Canonical, Reactome, Chemical/Genetic perturbations), and transcription factor motif gene sets. Additionally, several candidate gene sets were included from previous studies, including microarray analyses of HPr1-AR cells treated with DHT^7^, compiled gene sets previously utilized to encompass androgen response in PCa patients^13^, as well as gene sets differentiating basal from luminal prostate epithelial cells^90^. In total, 4204 gene sets were queried.

To identify meta-groups within enriched (NES > 1.8, FDR q-value < 0.05) gene sets, keywords from gene set identifiers were compiled, and frequency tables determined for all keywords across all gene sets and within enriched gene sets. To account for background frequencies of given terms across gene sets hypergeometric testing was used to determine if the frequency of key words within enriched gene sets was greater than expected.

### Validation of miR-96 target sites & Functional annotation

Unbiased assessment of miRNA seed sequence binding motifs in experimentally determined miR-96 target transcript 3’UTR regions was performed using the MSigDB microRNA targets collection and topGO package implemented in R. Significantly enriched terms (FDR < 0.05) by both methods were simultaneously visualized using Cytoscape 3.2.0.

## LIST OF ABBREVIATIONS

AR: Androgen receptor
CEBPB: CAAT/enhancer binding protein (C/EBP), beta,
ChIP: Chromatin Immunoprecipitation
CNV: Copy number variation
CTCF: CCCTC-binding factor (zinc finger protein)
DEGs: Differentially expressed genes
ENCODE: Encyclopedia of DNA elements
ERα: Estrogen receptor alpha
ETS: E26 transformation-specific
FAIRE: Formaldehyde-Assisted Isolation of Regulatory Elements
FDR: False discovery rate
GATA: GATA-BINDING PROTEIN 1
GEO: Gene Expression Omnibus
GSEA: Gene set enrichment analyses
H: Histone
HDAC: Histone deacetylase
K: Lysine
KDM1A/LSD1: Lysine (K)-Specific Demethylase 1A
miRNA: microRNA
NCOR1: Nuclear receptor corepressor 1
NCOR2/SMRT: Nuclear receptor corepressor 1/Silencing mediator for retinoid or thyroid-hormone receptors
NES: Normalized enrichment score
NR: Nuclear hormone receptor
ONEC2: Onecut2
PCa: Prostate cancer
PPAR: Peroxisome proliferator-activated receptor gamma
PPARGC1A: Peroxisome Proliferator-Activated Receptor Gamma, Coactivator 1 Alpha
PRAD: Prostate cancer cohort in TCGA
RARγ: Retinoic acid receptor gamma
RPKM: reads per kilobase per million mapped reads
Seq: Sequencing
shRNA: short hairpin RNA
SOX15: SRY (Sex Determining Region Y)-Box 15
STAT5A: Signal Transducer And Activator Of Transcription 5A
TACC1: Transforming, Acidic Coiled-Coil Containing Protein 1
TCGA: The cancer genome atlas
TNF: Tumor necrosis factor
TRIM28: Tripartite motif containing 28

## AUTHORS CONTRIBUTIONS

*MDL, PRvdB, MJC* all participated in the integrative analyses of RARγ functions. *MDL* generated and characterized the RARγ knockdown cells and all genomic data sets. *MDL, DJS, LESC, PKS, MDL* and *MJC* participated in study design and implementation. *MJC* conceived of the study, and participated in its design and coordination and directed the drafting the manuscript. All authors read and approved the final manuscript.

## ADDITIONAL FILES

17 supplemental figures are supplied.

One supplemental table is supplied.

